# Hippocampal and Parahippocampal Grey Matter Structural Integrity Assessed by Multimodal Imaging is Associated With Episodic Memory in Old Age

**DOI:** 10.1101/2020.02.07.936872

**Authors:** Ylva Köhncke, Sandra Düzel, Myriam C. Sander, Ulman Lindenberger, Simone Kühn, Andreas M. Brandmaier

## Abstract

Maintained structural integrity of hippocampal and cortical grey matter may explain why some older adults show rather preserved episodic memory. However, viable measurement models for estimating individual differences in grey matter structural integrity are lacking; instead, findings rely on fallible single indicators of integrity. Here, we introduce multitrait-multimethod (MTMM) methodology to capture individual differences in grey matter integrity, based on multimodal structural imaging in a large sample of 1,522 healthy adults aged 60 to 88 years from the Berlin Aging Study II, including 331 participants who underwent MR imaging. Structural integrity factors expressed the common variance of voxel-based morphometry (VBM), mean diffusivity (MD), and magnetization transfer ratio (MT) for each of four regions of interest (ROI): hippocampus, parahippocampal gyrus, prefrontal cortex, and precuneus. Except for precuneus, the integrity factors correlated with episodic memory. Associations with hippocampal and parahippocampal integrity persisted after controlling for age, sex, and education. Our results support the proposition that episodic memory ability in old age benefits from maintained structural integrity of hippocampus and parahippocampal gyrus. Exploratory follow-up analyses on sex differences showed that this effect is restricted to men. Multimodal factors of structural brain integrity might help improve our biological understanding of human memory aging.

---

Performance in episodic memory tasks typically declines after the age of 60 years (Rönnlund et al. 2005; Schaie et al. 1998), but there are pronounced age-related individual differences in levels and changes of performance (de Frias et al. 2007; Josefsson et al. 2012), with some older individuals displaying little or no performance decline. The “brain maintenance” hypothesis suggests that an older person’s level of behavioral performance reflects the degree to which this person’s brain has maintained its integrity across a variety of levels, including structure, function, and neurochemistry (Nyberg et al. 2012; Lindenberger 2014; Cabeza et al. 2018; Nyberg and Pudas 2019; Nyberg and Lindenberger in press). Modern neuroimaging techniques allow us to better describe and understand various characteristics of brain tissue through the application of different imaging modalities such as structural and functional magnetic resonance imaging (MRI), diffusion tensor imaging (DTI), or positron-emission tomography (PET). Yet, it is unclear whether and which of these measures converge on constructs that might reflect the “integrity” of a given brain region.

Here, we combine multimodal imaging with multitrait-multimethod (MTMM) modeling (Campbell and Fiske 1959; Eid et al. 2008) to represent the grey matter structural integrity of different regions of the human brain, and to investigate their associations with episodic memory. We selected a number of regions of interest (ROIs) that are part of the episodic memory network (for a review, see Dickerson and Eichenbaum 2009; Benoit and Schacter 2015). Specifically, we included hippocampus, parahippocampal gyrus, precuneus, dorsolateral prefrontal cortex, and medio-orbitofrontal cortex as ROIs. The hippocampus plays a key role in episodic memory (Eichenbaum 2017). Hippocampal volume is, on average, smaller in healthy older than in healthy younger adults, and shrinks with time in normal aging (Raz 2005; Raz, Lindenberger, et al. 2005; Fjell et al. 2009; Walhovd et al. 2011; Jäncke et al. 2019). Smaller hippocampal volume is related to poorer episodic memory performance in old age (for reviews, see Squire, 1992 or Kaup et al. 2011; Ward et al. 2015). Less decline in hippocampal volume in aging adults is related to less decline in episodic memory performance (J. Persson et al. 2012; Gorbach et al. 2017). Similar results were found at the functional level, such that smaller longitudinal decrements in activation were associated with better preservation of memory performance (J. Persson et al. 2012).

Regarding the role of hippocampal microstructural integrity in age-related cognitive decline, there is some indication that higher mean diffusivity (MD) in hippocampus, indicating a less dense tissue structure, is related to poorer episodic memory performance in older adults (Carlesimo et al. 2010). In studies using magnetization transfer imaging, it could be shown that a higher magnetization transfer ratio, indicating denser microstructure, is related to lower MD (Düzel et al. 2010), faster processing speed, and higher fluid intelligence (Aribisala et al. 2014), but not better memory (Düzel et al. 2010, 2008; Aribisala et al. 2014). Still, taken together, these findings suggest that the macro- and microstructural integrity of hippocampus might be critical for preserving its functionality for episodic memory in older age.

Hippocampus does not operate in isolation (Rugg et al. 2008; Dickerson and Eichenbaum 2009; Eichenbaum 2017). The parahippocampal gyrus is, together with the entorhinal cortex, a major input source for hippocampus, and critically supports episodic memory (J. Persson et al. 2012). Parahippocampal volume is smaller in older than in younger adults (Henson et al. 2016; Gorbach et al. 2017; Foster et al. 2019), and longitudinal decline in parahippocampal gyrus’ volume is related to decline in episodic memory performance (Gorbach et al. 2017), but cross-sectional associations between parahippocampal volume and episodic memory performance are not necessarily observed (Henson et al., 2016; Foster et al. 2019). MD in parahippocampal gyrus is higher in older than in younger adults (Grydeland et al. 2013).

Another region known to interact with hippocampus to support episodic memory is the prefrontal cortex (Eichenbaum 2017). Prefrontal cortex volume is shrinking with advancing age (Raz et al. 2005), its MD is higher in older than in younger individuals (Grydeland et al. 2013), and larger prefrontal grey matter volumes are related to better associative (Becker et al. 2015) as well as item episodic memory performance (N. Persson et al. 2017). Finally, precuneus contributes to memory retrieval (Cavanna and Trimble 2006), and MD in precuneus shows age-specific associations with cognitive performance (Grydeland et al. 2013).

*Brain maintenance* should be reflected by preserved tissue integrity on many biological levels, which most likely interact with one another as they change in the course of healthy aging. In structural imaging, brain maintenance should be reflected by relatively sparse and little microstructural damage as well as a relative lack of macrostructural atrophy. Maintenance can thus be assessed by several imaging modalities carried out at the same time in the same subjects. Such a multimodal imaging approach might provide a more comprehensive account of inter-individual variability in brain maintenance (Nyberg et al. 2012) than an approach that considers each measure separately. In the present study, we combine different structural characteristics of grey matter tissue into region-specific constructs of structural integrity. Our approach takes advantage of commonalities across measures from different modalities while removing the modality-specific measurement error; the rationale being that all selected measures reflect some aspect of structural integrity, so that their commonality should be a more robust index of structural integrity than any one measure alone.

Our approach differs from approaches adopted in other multimodal imaging studies. There are many good reasons to invest more effort into the use of multimodal brain imaging in aging research (Nevalainen et al. 2015; Fjell and Walhovd 2016). Many of the existing multimodal imaging studies aim at maximizing predictive accuracy by combining information from multiple modalities (Ritchie et al. 2015; Ward et al. 2015; Hedden et al. 2016; Liem et al. 2017). These approaches capitalize on the unique information that each modality adds to predicting cognitive performance (Ritchie et al. 2015; Ward et al. 2015; Hedden et al. 2016) or brain age (Liem et al. 2017), so they benefit from the fact that each modality measures different aspects of integrity. In contrast, in the present study, we were not primarily interested in the unique contribution of each measure to explaining an outcome, but instead in the variance that is shared across modalities. The presumed primary advantage of multimodality in our approach is that the resulting latent factor might yield a more reliable and valid estimate of the target concept, which is the purported tissue property of “structural integrity”. A latent factor of regional grey matter integrity expresses what the indicators have in common, is free of imaging modality-specific variance, and free of residual variance (measurement error). In comparison to currently available indicators, we postulate that a factor of this sort is more likely to do justice to the level of generality and abstraction that the term integrity suggests.

In this study, we aimed to quantify and jointly model different structural properties of grey matter by using three common imaging techniques, each being differentially sensitive to macro- and microstructural properties of brain tissue (Bartres-Faz and Arenaza-Urquijo 2011), namely grey matter volume, magnetization transfer (MT), and MD from DTI. In the next paragraph we describe which structural properties of grey matter are captured by the imaging modalities we selected for the current study.

Structural imaging provides static anatomical information derived from MR signal properties. T1-weighted, three-dimensional, high-resolution images are commonly used to estimate the volume of brain ROIs to study inter-individual differences in volume and volume changes over time. When using the voxel-based morphometry (VBM) method (Ashburner and Friston 2000; Good et al. 2001), signal intensity in every voxel is used to gauge regional variations in structural properties of the tissue and provides voxel-wise estimations of the local volume of specific tissue compartments (grey matter, white matter, or cerebrospinal fluid). Microstructural properties of grey matter regions, and age-related differences therein, can be probed by MT imaging (for a review, see Seiler et al. 2014). MT imaging capitalizes on the transfer of energy and related magnetization exchange between mobile water protons and protons that are immobilized by macromolecules (Wolff and Balaban 1989). MT ratio (MTR) values are calculated as the ratio between values measured with a MT pulse and values without MT pulse. MTR can detect subtle microstructural abnormalities due to age-related or pathological changes otherwise not detectable with standard MRI (Seiler et al. 2014). MTR values depend on content and concentration macromolecules bound to water molecules in relation to free water molecules. Lower MTR values can result from an increase in the mobile proton pool, occurring as a result of inflammation and edema, or a decrease in the semi-solid proton pool, associated with cell damage, axonal loss and demyelination (Seiler et al. 2014).

DTI can detect subtle changes in cellular microstructure by measuring patterns of water diffusion that likewise cannot be quantified using more traditional structural MRI sequences. MD is a DTI metric that measures the rate of water diffusion in all directions within an image voxel (Pierpaoli and Basser 1996) and is commonly used as an index of white matter microstructural integrity. MD can be used to characterize one form of structural integrity under the assumption that region-specific diffusion is based on (a) less diffusion across cell membranes in denser structures or structures with a main direction as seen in white matter tracts (Sundgren et al. 2004, Jespersen et al. 2007), (b) more diffusion within less dense brain structures or structures with no principal direction as seen in grey matter (Sundgren et al. 2004). Although more often used to characterize white matter, MD is also informative of grey matter microstructural properties and age-related differences in it (Abe et al. 2008; Grydeland et al. 2013), with lower MD indicating a denser structure, most probably indicating more cell membranes and intracortical myelin (Grydeland et al. 2013).

In the present study, we combined macro- and microstructural imaging modalities as indicators of grey matter integrity in a multimodal approach. Thus, we set out to validate a structural equation model representing the commonalities of specific tissue characteristics resulting from different imaging modalities. To establish the plausibility of integrity factors, we examined whether the empirically observed covariance structure shows substantial commonalities among the various grey-matter indices.

We used cross-sectional data from the older participants of the Berlin Aging Study II (BASE-II; Bertram et al. 2013), which amount to a fairly large sample of 1,532 healthy adults aged 60 to 88 years, with structural brain imaging measures of VBM, MTR, and MD taken from a subsample of 331 participants who underwent MR imaging. Our goal of this analysis approach was two-fold. First, we sought to demonstrate the benefits of a multivariate latent variable modeling approach to representing regional structural integrity while doing justice to the complexity of the underlying measurement problem. Second, we wished to demonstrate that such an approach can be put to use to identify the associations between structural properties of grey matter regions belonging to the episodic memory network and episodic memory ability in old age.

In a first set of analyses, we established region-specific latent brain integrity factors. To this end, we specified confirmatory factor models within each of the brain regions by defining a latent brain integrity factor representing the variance that is shared across the three imaging modalities. This latent factor should capture the statistical communality of different physical properties of grey matter tissue. This parallels psychometric approaches targeting a non-observable (or latent) psychological construct by measuring a range of indicators and interpreting their common variance as representative of the target construct. By using different indicators, we triangulate our target construct, integrity, which is sensible even if our indicators should only have limited overlap in variance (Little et al. 1999).

In a second set of analyses, we combined the latent brain integrity model with a latent episodic memory factor to investigate the associations of brain integrity and episodic memory performance. We hypothesize that grey matter integrity in regions of the episodic memory network is related to episodic memory performance.

## Methods

### Participants and Study Design

Healthy older participants were recruited from BASE-II, a multi-institutional and multidisciplinary study assessing variables from a wide range of domains for each participant (Bertram et al. 2013). Participants completed a comprehensive cognitive examination (see Düzel et al. 2016, for further details). A subsample of eligible participants was then invited to take part in a separate MRI session within a couple of weeks (mean time interval 3.8 months, SD = 4.4) after completing cognitive testing. None of the participants took any medication that is known to potentially affect memory function or had a history of head injuries, medical (e.g., heart attack), neurological (e.g., epilepsy), or psychiatric disorders (e.g., depression). Additionally, all participants had completed at least 8 years of education. BASE-II includes a larger sample of persons above age 60 and a smaller sample of participants in early adulthood. Here, we selected only data from participants above age 60, of which 1532 had completed cognitive testing, of which 340 had additionally taken part in MR imaging. We had to exclude 9 cases with erroneous data from the cognition sample, 2 of which were in the MR sample as well. We then excluded multivariate outliers with highly unlikely combinations of values (*p* < .0001 of robust Mahalanobis distances; detected using R-package faoutlier, version 0.7.2, Chalmers and Flora 2015, method “mve”, in complete cases only). We detected multivariate outliers separately for the 4 episodic memory tests in the total sample (n = 1523; 1 outlier found) and for the 12 MR variables in the MR sample (n = 338; 7 outliers found). Hence, the effective sample with cognitive data consisted of 1522 older adults (Table 1), the effective sample with MR-data consisted of 331 older adults (Table 2). The ethics committee of the Max Planck Institute for Human Development had approved the cognitive testing and the ethics committee of the Deutsche Gesellschaft für Psychologie (DPGs) had approved the imaging study. Participants received monetary compensation for their participation in the cognitive and imaging sessions and provided informed consent in accordance to the Declaration of Helsinki.

**Table 1.** The descriptive statistics of the background variables and episodic memory task scores both in the full sample and the MR sample, and selectivity (in SD) of the MR sample.

| Variable | N | Mean |  | SD |  | Skewness |  | Kurtosis |  | Selectivity<sup>1</sup> |
| --- | --- | --- | --- | --- | --- | --- | --- | --- | --- | --- |
|  |  | Total sample | MR sample | Total sample | MR sample | Total sample | MR sample | Total sample | MR sample |  |
| **Age** | 1511 | 70.5985 | 70.1892 | 3.8381 | 3.8858 | 0.3127 | 0.2898 | 3.5882 | 3.0102 | 0.1066 |
| **Sex (% f)** | 1522 | 50.72 | 38.07 |  |  |  |  |  |  |  |
| **Education** | 1339 | 14.1621 | 14.0845 | 2.8915 | 2.8874 | 0.1385 | 0.1267 | 1.6151 | 1.6937 | 0.0268 |
| **VLMT** | 1497 | 8.4963 | 8.445 | 2.6809 | 2.5422 | -0.4279 | -0.3522 | 2.4881 | 2.6238 | 0.0191 |
| **FP** | 1495 | 0.2665 | 0.2729 | 0.2097 | 0.1992 | -0.0217 | 0.1139 | 2.8945 | 2.7007 | -0.0306 |
| **SE** | 1495 | 0.2776 | 0.2906 | 0.1397 | 0.1352 | -0.0215 | -0.1097 | 2.9302 | 2.9105 | -0.093 |
| **OL** | 1505 | 13.2551 | 13.5583 | 4.0043 | 4.0415 | 0.0478 | 0.0969 | 3.1182 | 3.4811 | -0.0757 |

*Note.* <sup>1</sup> Selectivity = $(\text{mean}_{(\text{total sample})} - \text{mean}_{(\text{MR sample})}) / \text{SD}_{(\text{total sample})}$ , VLMT = Verbal Learning and Memory Test – sum of remembered words across 5 learning trials (all with the same 15 words), FP = Face–Profession task – hits minus false alarms, SE = Scene Encoding task – hits minus false alarms, OL = Object Location task – sum of correct placements across 2 trials (each consisting of 12 items).

**Table 2.**
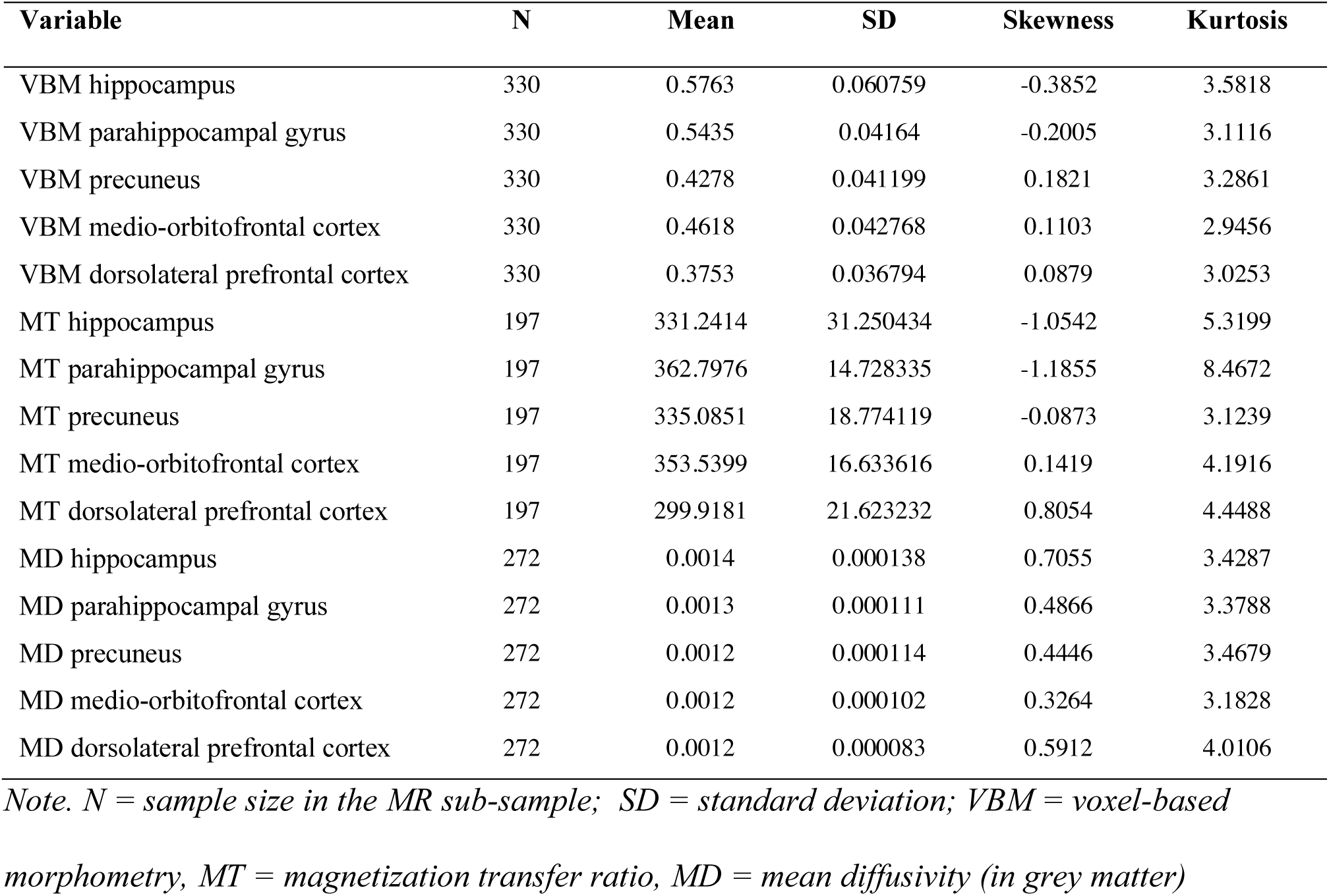
Descriptive statistics of the MR variables, in the MR sample

| Variable | N | Mean | SD | Skewness | Kurtosis |
| --- | --- | --- | --- | --- | --- |
| VBM hippocampus | 330 | 0.5763 | 0.060759 | -0.3852 | 3.5818 |
| VBM parahippocampal gyrus | 330 | 0.5435 | 0.04164 | -0.2005 | 3.1116 |
| VBM precuneus | 330 | 0.4278 | 0.041199 | 0.1821 | 3.2861 |
| VBM medio-orbitofrontal cortex | 330 | 0.4618 | 0.042768 | 0.1103 | 2.9456 |
| VBM dorsolateral prefrontal cortex | 330 | 0.3753 | 0.036794 | 0.0879 | 3.0253 |
| MT hippocampus | 197 | 331.2414 | 31.250434 | -1.0542 | 5.3199 |
| MT parahippocampal gyrus | 197 | 362.7976 | 14.728335 | -1.1855 | 8.4672 |
| MT precuneus | 197 | 335.0851 | 18.774119 | -0.0873 | 3.1239 |
| MT medio-orbitofrontal cortex | 197 | 353.5399 | 16.633616 | 0.1419 | 4.1916 |
| MT dorsolateral prefrontal cortex | 197 | 299.9181 | 21.623232 | 0.8054 | 4.4488 |
| MD hippocampus | 272 | 0.0014 | 0.000138 | 0.7055 | 3.4287 |
| MD parahippocampal gyrus | 272 | 0.0013 | 0.000111 | 0.4866 | 3.3788 |
| MD precuneus | 272 | 0.0012 | 0.000114 | 0.4446 | 3.4679 |
| MD medio-orbitofrontal cortex | 272 | 0.0012 | 0.000102 | 0.3264 | 3.1828 |
| MD dorsolateral prefrontal cortex | 272 | 0.0012 | 0.000083 | 0.5912 | 4.0106 |

*Note.* *N* = sample size in the MR sub-sample; *SD* = standard deviation; *VBM* = voxel-based
*morphometry*, *MT* = magnetization transfer ratio, *MD* = mean diffusivity (in grey matter)

### MRI Acquisition

Images were acquired on a Siemens Tim Trio 3T scanner (Erlangen, Germany) using a 32-channel head coil. The T1 images were obtained using a three-dimensional T1-weighted magnetization prepared gradient-echo (MPRAGE) sequence based on the ADNI protocol (www.adni-info.org; repetition time (TR) = 2500 ms; echo time (TE) = 4.77 ms; TI = 1100 ms, acquisition matrix = 256 × 256 × 176, flip angle = 7°; 1 x 1 x 1 mm^3^ voxel size). Diffusion-weighted images were obtained with a single-shot diffusion-weighted spin-echo-refocused echo-planar imaging sequence (FOV 218 mm × 218 mm; 128 × 128 matrix interpolated to 256 × 256; TE = 98 ms; TR = 11000 ms; 73 slices; slice thickness 1.7 mm; b-value 1000 s/mm2; 60 directions). MT images consisting of two volumes were acquired with identical settings (transversal, 256×256 pixels, TE = 5.5 ms, TR = 28ms 48 slices, voxel size 1mm×1mm×3mm). The first image (MT image) was acquired with a magnetic saturation pulse (1200 Hz off-resonance, 16 ms) and the second (noMT image) without a magnetic saturation pulse resulting in a proton-density-like image.

### MR Preprocessing

#### Voxel-based morphometry (VBM)

Structural data were processed with CAT12 (Computational Anatomy Toolbox 12, Structural Brain Mapping group, Jena University Hospital, Jena, Germany), a toolbox that is implemented in SPM12 (Statistical Parametric Mapping, Institute of Neurology, London, UK) for voxel-based morphometry analysis of imaging data. We applied the CAT12 default cross-sectional pre-processing stream, which implements correction of the T1-weighted images for bias-field inhomogeneities, segmentation into grey matter, white matter and CSF, and spatial normalized using the DARTEL (diffeomorphic anatomical registration through exponentiated lie algebra) algorithm. Modulation with Jacobian determinants was applied in order to preserve the volume of a particular tissue within a voxel leading to a measure of volume of grey matter. Grey matter images were used for the current set of analyses and smoothed with a Gaussian kernel of 8 mm (full width at half maximum).

#### Magnetization transfer imaging (MTI)

The MTR maps for each subject were calculated on a voxel-by-voxel basis according to the formula MTR = (noMT-MT) / noMT. The data were then normalized into MNI space.

#### Diffusion tensor imaging (DTI)

Diffusion-weighted images were pre-processed using the FSL software package (Smith et al. 2004; Jenkinson et al. 2012), version 5.0. This included corrections of potential head movement and inspection of image quality. The first non-diffusion weighted image of each individual image set was used as a brain mask. The difference in alignment between this initial image and recurrent ones in the sequence was estimated using FLIRT (FMRIB’s Linear Image Registration Tool; Jenkinson et al. 2002) and then corrected for by means of re-alignment. The resulting data was then processed via FSL’s *dtifit* to fit a diffusion tensor model at each voxel and obtain the MD values. The MNI based maps were produced using the standard TBSS pipeline (Smith et al. 2006).

### ROI Extraction

Based on prior studies on associations between regional grey matter structure and episodic memory (Becker et al. 2015; Cavanna and Trimble 2006; Gorbach et al. 2017; Pantel et al. 2003; Raz and Rodrigue 2006; Squire 1992) as well as functional correlates of episodic memory (Benoit and Schacter, 2015 J. Persson et al. 2012), we extracted ROIs bilaterally from the hippocampus, medio-orbitofrontal cortex, dorsolateral prefrontal cortex, parahippocampal gyrus, and precuneus, as defined by the Automated Anatomical Labelling (AAL) atlas (Tzourio-Mazoyer et al. 2002).

### Episodic Memory Assessment

All BASE-II participants were invited to two cognitive test sessions with an exact interval of seven days and at the same time of day to avoid circadian confounding effects on performance. Each session lasted about 3.5 h. Participants were tested in groups of 4–6 individuals. Each group was instructed via a standardized session manual. Each task started with a practice trial to ensure that every participant understood the task. Depending on the task, responses were given via button boxes, the computer mouse, or a keyboard.

The cognitive battery of BASE-II covers key cognitive abilities measured by 21 tasks, four of which assess aspects of episodic memory and were thus selected for the present study: (i) *Verbal Learning and Memory*, assessing free recall of auditorily presented words after each of five learning trials (mean across 5 learning trials); (ii) *Face–Profession,* testing associative recognition memory 5 minutes after incidental encoding; (iii) *Scene Encoding,* measuring recognition memory of incidentally encoded scenes after a delay of 2.5 hours; (iv) *Object Location,* assessing free recall of deliberately encoded object locations. The tasks are described in detail in Düzel et al. (2016) and in the supplementary materials. As reported before (Düzel et al. 2016; Kühn et al. 2017), performance on these four tasks is well captured by a latent factor of episodic memory ability. A one-factor model fits the data well both in the total sample, CFI > .999; RMSEA < .001; SRMR = .004, and in the MR sample, CFI > .999; RMSEA < .001; SRMR = .017.

### Statistical Analyses

We used structural equation modeling to investigate the relationship between episodic memory performance and structural grey matter integrity in a multivariate approach, for two reasons. First, it enables us to capture variance shared across three different structural brain-imaging modalities in a latent factor of structural integrity for any given brain region. This makes sense from a theoretical perspective, as we aim to define a statistically plausible index of region-specific grey matter structural integrity. While the three imaging modalities are designed to assess different characteristics of grey matter structure, their shared variance can be interpreted as indicating a common cause for relatively good or relatively poor grey matter structural integrity. By separating the shared variance (i.e., what is common across measures) from the unique, modality-specific variance (i.e., what is specific to the measurement instrument), we hope to acquire a more reliable and valid estimate of integrity. In addition, we defined latent method factors for each modality (VBM, MTR, and MD) that capture common variation across the modality across all regions (capturing what is common to the measurement instrument only but not to the common factor). As a consequence, the residual variance estimates in our model represent variance that is neither shared by all ROIs in a given modality nor shared by all modalities in a given ROI (e.g., measurement error).

Second, we used structural equation modelling to examine whether the region-specific factors of grey matter structural integrity were associated to episodic memory performance. In this context, a particular virtue of structural equation modelling is that we can model grey matter integrity for each of the ROIs as well as episodic memory performance as latent factors.

We specified and estimated structural equation models in Onyx (Oertzen et al. 2014), version 1.0-1029, and lavaan (Rosseel 2012), version 0.6-5, a SEM package in R (R Core Team 2016), version 3.6.2 (2019-12-12). To account for missing data, we used full information maximum likelihood estimation. Given that large differences in measurement scales, like those in our data, typically pose problems for numerical optimization algorithms, all observed variables were rescaled to a mean of 5 and standard deviation of 2. To evaluate model fit, we used the Root Mean Square Error of Approximation (RMSEA), the Comparative Fit Index (CFI), and the Standardized Root Mean Residual (SRMR) (Schermelleh-Engel et al. 2003). We interpret an RMSEA < .08, a CFI > .90, and SRMR < .8 as acceptable model fit. To assess statistical significance of parameters, we used the likelihood ratio test, that is, we compared a model with the parameter of interest freely estimated to a nested model with this parameter fixed to zero, and compared whether the χ^2^ difference between the models indicated a significant difference in fit (Kline 2011). For loadings and variance parameters, we used Wald tests.

In a first set of analyses, we specified separate CFAs to validate each latent integrity factor model for the selected ROIs (dorsolateral prefrontal cortex, medio-orbitofrontal cortex, hippocampus, parahippocampal gyrus, precuneus) defined by the indicators representing the three imaging modalities. In a next set of analyses, we combined the individual, fully saturated models to one structural equation model, in which all latent integrity factors were allowed to covary with one another. The intercorrelation of the medio-orbitofrontal cortex factor and the dorsolateral prefrontal cortex factor was too high for the two factors to be meaningfully modeled as separate latent variables (estimated r > 1), hence we decided to specify a single latent prefrontal cortex factor with six indicators, two per modality, from medio-orbitofrontal cortex and dorsolateral prefrontal cortex. Next, we added modality-specific latent factors (methods factors), which were defined to be orthogonal to the ROI factors, so that they represent the modality-specific share of the variance in the measures after the ROI-specific variance is accounted for. The methods factors were allowed to correlate with one another, and to be measured by all indicators that were derived from the respective modality, with loadings freely estimated (see Figure 3). This type of model is known as a MTMM (multi-trait multi-method) model (Campbell and Fiske 1959; Eid et al. 2008). It is the appropriate measurement model if multiple characteristics (usually traits, but here, ROIs) are each measured by several distinct measures (usually raters, but here, imaging modalities), yielding a latent integrity factor for each ROI and a latent method factor for each imaging modality. The latent scales of both the ROI factors and method factors were identified by fixing the loading of a reference indicator to one, which was the respective VBM indicator for the ROI factors and the respective measure of precuneus for the method factors. Within this model, we then correlated the ROI factors and the method factors with age to investigate age differences in variables of interest. We expected age differences for all ROI integrity factors (Fjell et al. 2009; Grydeland et al. 2013; Raz et al. 2005; Seiler et al. 2014). We did not formulate any specific hypotheses for age differences in the methods factors, because method-specific variance was not in the focus of the investigation. Age differences in any method factor would indicate that this method is especially age-sensitive, over and above the age-related variance it shares with the other methods and that is captured in the ROI-wise integrity factors. Another purpose of investigating age differences was to be able to statistically control for potential age effects on the brain–behavior associations.

The first set of analyses served to establish the latent integrity factors. In a second set of analyses, we investigated associations between grey matter structure and episodic memory. We added a latent factor of episodic memory performance to the model that we defined by the four indicators of the episodic memory tasks described earlier (Düzel et al. 2016; Kühn et al. 2017), and estimated associations between the latent episodic memory factor and the ROI integrity factors. We set up the latent associations in two ways that give alternative interpretations but are statistically equivalent: once as covariances to estimate the first-order correlations among ROIs and episodic memory (correlational model) and once as regressions to assess the unique associations of each ROI factor with episodic memory while controlling for the other factors’ association with it (regression model). With the regression model, we were able to assess how much variance in episodic memory each ROI factor accounts for, over and above the other ROI factors. In this way, we sought to disentangle general associations of any brain region’s grey matter structure from associations that are unique to a specific brain region’s grey matter structure, given the ROIs represented in the model. In addition, we estimated the total variance in episodic memory that all ROI factors together accounted for (see MIMIC model in Kievit et al. 2012, p 93).

In a next step, we entered age and education (years) as well as sex into the correlational model to statistically control for the extent to which potential associations between episodic memory and grey matter being are being caused by these covariates. With respect to age, we expected that older participants tended to show lower grey matter integrity and lower episodic memory ability, so that not controlling for age in this age-heterogeneous sample would most likely yield a strong association between grey matter and episodic memory that is at least partly driven by those age differences. Education was expected to be related to episodic memory performance such that persons with more years of education tend to score better in episodic memory tests (Stern 2009). Education may affect performance as a consequence from education-related early training of memory, but also serves as a proxy for socio-economic status, differences in which are not of interest in this study. Sex differences in episodic memory performance were to be expected (for a review, see Asperholm et al. 2019), and possibly even in grey matter structure (for a review, see Ruigrok et al. 2014). These could then could induce an association if not adjusted for. We even explored sex differences in the association structures (see supplementary material).

As an additional ad-hoc exploratory analysis, we investigated differences between men and women in the factor models and in the associations between grey matter integrity and episodic memory (supplementary material).

Models that entailed only brain data or brain data and age were fitted to data from the MR sample (n = 331), and models that entailed episodic memory data were fitted to data from the total sample, under the assumption that the MR data were missing at random (Rubin 1976; Schafer and Graham 2002). This assumption holds as long as missingness in the grey matter variables is either completely random or can be explained by variables in the model.

## Results

### Sample Descriptives

Table 1 displays the descriptive statistics of the selected covariates and episodic memory task performance variables in the full sample and the MR sample. None of the variables were heavily skewed, and kurtosis was high only in the Verbal Learning and Memory test in the full sample, so we assumed that all variables follow the normal distribution to an acceptable extent. The MR sample did not differ much from the full BASE-II sample, with selectivity below 0.11 standard deviations in the continuous variables. The only considerable difference was found in the sex distribution, with about 38% females in the MR sample and 51% females in the full sample. In Table 2, we report the descriptive statistics of the grey matter structure variables in their original scale in the MR sample. We deemed skewness and kurtosis levels acceptable in these variables, too. Days between the cognitive assessment and the MR-session differed between individuals (absolute difference in days: mean = 113.5, SD = 125.6, min = 8, max = 774). For sex differences in all variables of interest, see the supplementary material. For pairwise correlations between all variables of interest see Table 3.

**Table 3.** Pairwise first-order correlations between all observed variables of interest

|  |  | Memory |  |  |  | VBM |  |  |  | MT |  |  |  | MD |  |  |  | Covariates |  |  |  |  |
| --- | --- | --- | --- | --- | --- | --- | --- | --- | --- | --- | --- | --- | --- | --- | --- | --- | --- | --- | --- | --- | --- | --- |
|  |  | VL MT | FP | SE | OL | HC | PHG | PRE | OFC | DL PFC | HC | PHG | PRE | OFC | DL PFC | HC | PHG | PRE | OFC | DL PFC | age | edu |
| Memory | FP | 0.25* |  |  |  |  |  |  |  |  |  |  |  |  |  |  |  |  |  |  |  |  |
|  | SE | 0.26* | 0.22* |  |  |  |  |  |  |  |  |  |  |  |  |  |  |  |  |  |  |  |
|  | OL | 0.30* | 0.25* | 0.29* |  |  |  |  |  |  |  |  |  |  |  |  |  |  |  |  |  |  |
| VBM | HC | 0.12* | 0.15* | 0.14* | 0.14* |  |  |  |  |  |  |  |  |  |  |  |  |  |  |  |  |  |
|  | PHG | 0.05 | 0.10 | 0.00 | -0.03 | 0.68* |  |  |  |  |  |  |  |  |  |  |  |  |  |  |  |  |
|  | PRE | 0.02 | 0.08 | 0.05 | 0.06 | 0.27* | 0.30* |  |  |  |  |  |  |  |  |  |  |  |  |  |  |  |
|  | OFC | 0.09 | 0.09 | 0.09 | 0.06 | 0.32* | 0.31* | 0.38* |  |  |  |  |  |  |  |  |  |  |  |  |  |  |
|  | DLPFC | 0.10 | 0.09 | 0.06 | 0.03 | 0.34* | 0.34* | 0.44* | 0.58* |  |  |  |  |  |  |  |  |  |  |  |  |  |
| MT | HC | 0.16* | 0.24* | 0.21* | 0.06 | 0.68* | 0.32* | 0.17* | 0.37* | 0.25* |  |  |  |  |  |  |  |  |  |  |  |  |
|  | PHG | 0.17* | 0.18* | 0.13 | -0.04 | 0.42* | 0.29* | 0.16* | 0.34* | 0.28* | 0.69* |  |  |  |  |  |  |  |  |  |  |  |
|  | PRE | 0.07 | 0.08 | 0.08 | 0.01 | 0.18* | 0.16* | 0.55* | 0.12 | 0.16* | 0.18* | 0.32* |  |  |  |  |  |  |  |  |  |  |
|  | OFC | 0.06 | 0.10 | 0.07 | 0.01 | 0.33* | 0.13 | 0.07 | 0.28* | 0.26* | 0.46* | 0.45* | 0.28* |  |  |  |  |  |  |  |  |  |
|  | DLPFC | -0.03 | 0.08 | 0.02 | -0.08 | 0.21* | 0.08 | 0.16* | 0.23* | 0.33* | 0.31* | 0.34* | 0.52* | 0.57* |  |  |  |  |  |  |  |  |
| MD | HC | -0.05 | -0.16* | -0.26* | -0.12* | -0.64* | -0.27* | -0.14* | -0.28* | -0.18* | -0.86* | -0.45* | -0.12 | -0.33* | -0.25* |  |  |  |  |  |  |  |
|  | PHG | -0.14* | -0.13* | -0.26* | -0.10 | -0.56* | -0.43* | -0.40* | -0.41* | -0.39* | -0.68* | -0.62* | -0.38* | -0.35* | -0.31* | 0.64* |  |  |  |  |  |  |
|  | PRE | -0.09 | -0.12* | -0.16* | -0.07 | -0.32* | -0.29* | -0.64* | -0.25* | -0.31* | -0.26* | -0.38* | -0.76* | -0.30* | -0.36* | 0.29* | 0.57* |  |  |  |  |  |
|  | OFC | -0.06 | -0.13* | -0.13* | -0.04 | -0.25* | -0.25* | -0.40* | -0.32* | -0.39* | -0.25* | -0.27* | -0.46* | -0.46* | -0.60* | 0.23* | 0.51* | 0.65* |  |  |  |  |
|  | DLPFC | -0.12* | -0.14* | -0.19* | -0.03 | -0.40* | -0.33* | -0.31* | -0.43* | -0.40* | -0.47* | -0.50* | -0.40* | -0.65* | -0.54* | 0.47* | 0.65* | 0.55* | 0.71* |  |  |  |
| Covariates | age | -0.13* | -0.16* | -0.16* | -0.05 | -0.28* | -0.19* | -0.11* | -0.12* | -0.17* | -0.26* | -0.27* | -0.20* | -0.29* | -0.23* | 0.32* | 0.38* | 0.30* | 0.41* | 0.46* |  |  |
|  | edu | 0.18* | 0.16* | 0.09* | 0.15* | -0.10 | -0.06 | -0.04 | 0.00 | -0.02 | -0.01 | -0.04 | -0.11 | 0.08 | -0.03 | 0.05 | 0.10 | 0.11 | -0.01 | 0.00 | -0.06* |  |
|  | sex | 0.09* | 0.03 | 0.04 | 0.04 | 0.44* | 0.26* | 0.30* | 0.29* | 0.33* | 0.38* | 0.28* | 0.27* | 0.27* | 0.17* | -0.27* | -0.41* | -0.29* | -0.18* | -0.31* | 0.02 | -0.15* |

Note. VLMT = Verbal Learning and Memory test; FP = Face-Profession task; SE = Scene Encoding; OL = Object Location task; V = VBM (volume-based morphometry); MT = Magnetization transfer ratio; MD = mean diffusivity; HC = hippocampus; PHG = parahippocampal gyrus; PRE = precuneus; MOFC = medio-orbitofrontal cortex; DLPFC = dorsolateral prefrontal cortex; edu = years of education. Based on pairwise complete data. \* $p < .05$ .

### Model of Grey Matter Integrity

We succeeded in establishing a factor model with four ROI factors capturing shared variance across VBM, MT, and MD within each ROI (prefrontal cortex, hippocampus, parahippocampal gyrus, precuneus). We extracted ROI-wise data from 5 ROIs, but 2 prefrontal ROIs shared such a large part of their variance that they were equally well represented by a common latent factor (prefrontal cortex). In addition, the model included method-specific factors (V, MT, MD) that were orthogonal to the ROI factors and captured the shared variance of measures within imaging modalities and across ROIs (Figure 2). We allowed for residual covariances between the VBM indicators of closely neighboring ROIs (namely, of medio-orbitofrontal and dorsolateral prefrontal cortex and of hippocampus and parahippocampal gyrus), as we expected dependencies between them to exceed the shared variance modeled in the latent V factor. Fixing these residual covariances to zero (as done with all other residual covariances among indicator variables) significantly decreased model fit. The proposed model with the residual covariances fitted the data well, *χ^2^* (df=64) = 136.05; *CFI* = .972; *RMSEA* = .058; *SRMR* = .04. All observed variables loaded reliably on the postulated latent ROI factors except for MD of medio-orbitofrontal cortex and MT of dorsolateral prefrontal cortex on the prefrontal factor (standardized loadings = 0.06 and 0.1, *z*’s = .053 and 1.05, *p*’s = 0.6 and 0.29, all other standardized loadings > 0.14, *z*’s > 2.27, *p*’s < 0.023). All indicators loaded reliably on the postulated method factors (standardized loadings > .25 and > .94). We estimated covariances among the ROI integrity factors and among method factors, while method factors were defined as being orthogonal to ROI factors. Covariances among the ROI integrity factors were all significant except for the one between prefrontal cortex and precuneus (see Table 4). The covariances among the method factors were non-significant except for the one between VBM and MD (Table 4).

**Table 4.** Correlations among latent factors in the grey matter integrity model

|  | *Prefrontal cortex* | *Hippocampus* | *Parahippocampal g.* | *Precuneus* | *Method VBM* | *Method MD* | *Method MT* |
| --- | --- | --- | --- | --- | --- | --- | --- |
| *Prefrontal cortex* | 1 |  |  |  |  |  |  |
| *Hippocampus* | .58 <sup>a)</sup> | 1 |  |  |  |  |  |
| *Parahippocampal g.* | .61* | .82* | 1 |  |  |  |  |
| *Precuneus* | -.09 | .02* | .24* | 1 |  |  |  |
| *Method: VBM* | 0 <sup>b)</sup> | 0 <sup>b)</sup> | 0 <sup>b)</sup> | 0 <sup>b)</sup> | 1 |  |  |
| *Method: MD* | 0 <sup>b)</sup> | 0 <sup>b)</sup> | 0 <sup>b)</sup> | 0 <sup>b)</sup> | -.66* | 1 |  |
| *Method: MT* | 0 <sup>b)</sup> | 0 <sup>b)</sup> | 0 <sup>b)</sup> | 0 <sup>b)</sup> | .47* | -.82* | 1 |

Note. \* $p < .05$ ; according to $\chi^2$ difference test / likelihood ratio test with 1 df. <sup>a)</sup> Model with this covariance restricted to 0 did not converge, so the Wald test was used instead for statistical inference. <sup>b)</sup> The parameter was defined as zero in the model.

### Association Between Grey Matter Integrity Factors and Age

We included age in the model to estimate covariances between age and all latent factors. Model fit remained acceptable, *χ^2^_(df=72)_* = 148.66*; CFI* = .97*; RMSEA* = .057*; SRMR* = .038. Age was negatively associated with all ROI factors except precuneus, *r_age, PFC_* = −.20, Δ *χ^2^_(df = 1)_* = 6.9, *p* = .0085; *r_age, HC_* = −.21, Δ *χ^2^_(df = 1)_* = 13.5, *p* = .0002; *r_age, PHG_* = −.20, Δ *χ^2^_(df = 1)_ =* 8.4, *p* = .0038; *r_age, PRE_* = .004, Δ *χ^2^_(df = 1)_* = .004; *p* = .95. In addition, age was associated with the method factors (*r_age, V_* = −.18, Δ *χ^2^_(df = 1)_* = 7.04, *p* = .008; *r_age, MT_* = −.32, Δ *χ^2^_(df = 1)_* = 19.8, *p* < .0001; *r_age, MD_* = .43, Δ *χ^2^_(df = 1)_* = 46.4, *p* < .0001).

### Associations Between Grey Matter Integrity ROIs and Episodic Memory in the Correlational Model

We added an episodic memory latent factor to the model, which was indexed by the four episodic memory task scores, and estimated correlations between the episodic memory factor and the ROI integrity factors (correlational model). We fit this model to the data from all included participants with episodic memory data available (*n* = 1,522), assuming missingness of MR data to be at random. The fit was acceptable, *χ^2^_(df =119)_* = 190.17*; CFI* = .98*; RMSEA* = .02*; SRMR* = .04. The scores that served as indicators all loaded significantly on the latent episodic memory factor with moderate effect size (standardized loadings between .46 and .55, *z*’s > 10.41, *p*’s < .001), indicating that they contributed similarly to the latent factor. Episodic memory was significantly associated with all ROIs except precuneus, *r _EM, PFC_* = .23, Δ *χ^2^_(df = 1)_ = 5.08*; *p* = .024; *r _EM, HC_* = .33, Δ *χ^2^_(df = 1)_ = 16.2*; *p* < .0001; *r _EM, PHG_* = .30, Δ *χ^2^_(df = 1)_* =14.04; *p* = .0013; *r _EM, PRE_* = .08, Δ *χ^2^_(df = 1)_ = 0.64*; *p* = .42. In contrast, episodic memory was not reliably related to the method factors, *r _EM, V_* = .10, Δ *χ^2^_(df = 1)_ = 1.17, p* = .28; *r _EM, MT_* = .03, Δ *χ^2^_(df = 1)_ = .08*; *p* = .77; *r _EM, MD_* = −.12, Δ *χ^2^_(df = 1)_ = 1.96*; *p* = .16. Hence, a model with these associations fixed to zero also had a good fit to the data, *χ^2^ (df=122)* = 192.56*; CFI* = .98*; RMSEA* = .02*; SRMR* = .04, and was used in the following extensions of the model.

### Unique Associations of ROIs With Episodic Memory in Regression Model

To assess how much variance in episodic memory is uniquely and jointly predicted by the latent ROI factors, we refit the previous model with directed paths from each ROI to latent episodic memory (regression model). This is similar to a latent multiple regression model regressing episodic memory on each ROI integrity factor. Note that the directionality of effects (i.e., ROI integrity factor predicting memory) is merely hypothesized and cannot be tested with the data at hand. Importantly, instead of interpreting first-order correlations, we now examined the total effect and the unique effects of each ROI integrity factor on episodic memory. All ROIs together explained 12.9% of the variance in episodic memory. No ROI predicted episodic memory over and above its shared variance with the other ROIs. However, hippocampus had the largest effect size; it was greater than precuneus, parahippocampal gyrus, and prefrontal cortex by a factor of 3, 3.4, and 4, respectively (*std.β _EM,HC_* = .24, Δ *χ^2^_(df = 1)_ = 1.48*; *p* = .22; *std.β _EM,PRE_* = .08, Δ *χ^2^_(df = 1)_ = .48*; *p* = .49; *std.β _EM,PHG_* = .07, Δ *χ^2^_(df = 1)_* = .08; *p* = .77; *std.β _EM,PFC_* = .06, Δ *χ^2^_(df = 1)_ = .18*; *p* = .68).

**Figure 1.**
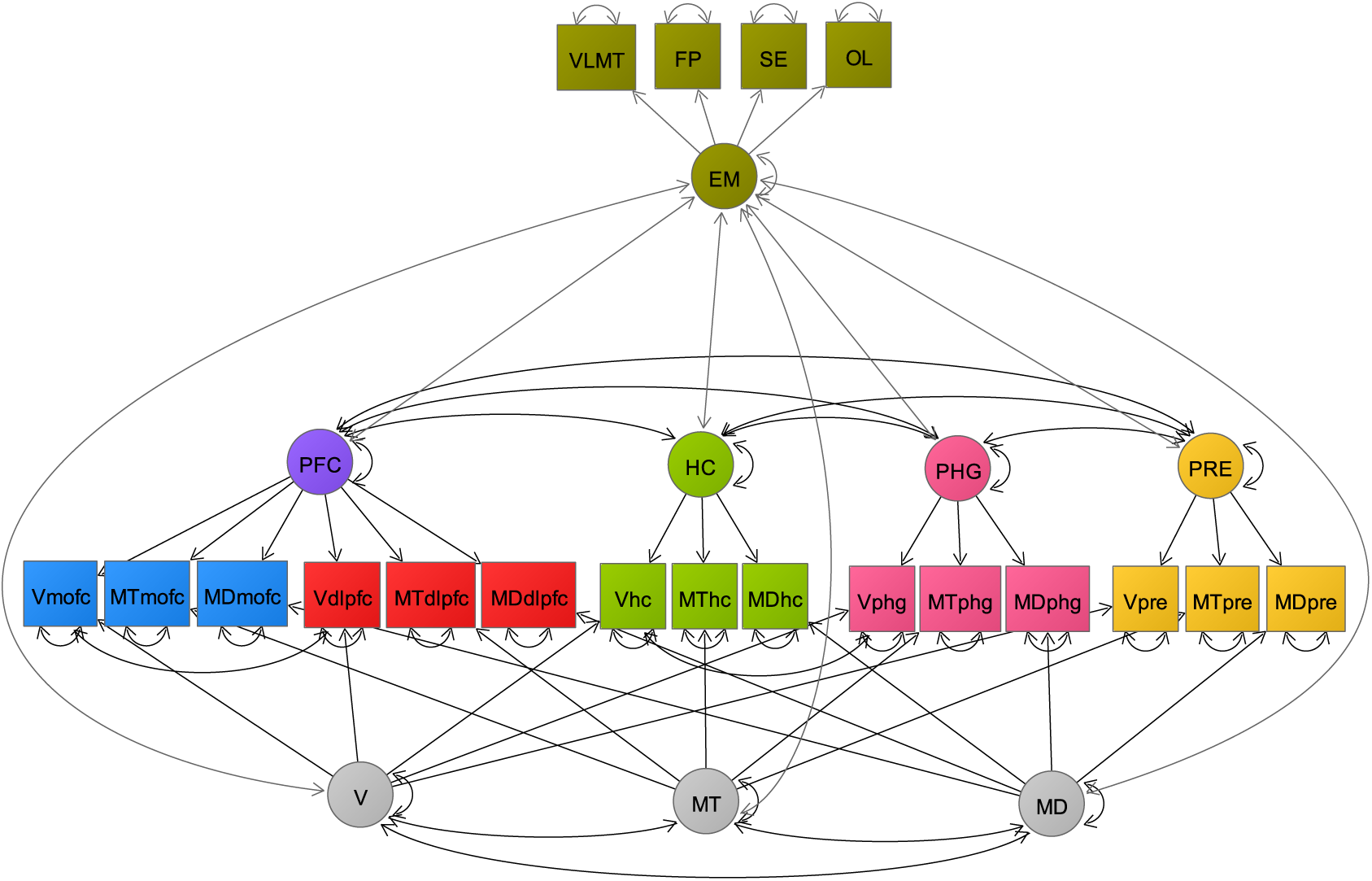
Correlation model. Note. Grey matter integrity factor model separating common variance across structural imaging modalities for each of the ROIs (PFC, HC, PHG, PRE) from method-specific variance (V, MT, MD) and residual variance (double-headed arrows at the observed variables depicted by squares), and associating grey matter factors with episodic memory. Circles depict latent variables, squares depict observed variables. Double-headed arrows are variances or covariances, single-headed arrows are regression parameters or loading parameters. Names beginning with V are VBM-derived grey matter probability measures, names with MT are magnetization transfer ratio measures, names with MD mean diffusivity measures, further, names indicate the regions; mofc = medio-orbitofrontal cortex, dlpfc = dorsolateral prefrontal cortex, hc = hippocampus; phg = parahippocampal gyrus. Upper part in light grey: Episodic memory factor added in correlation model. EM = episodic memory, VLMT = verbal learning and memory test, FP = face-profession task, SE = scene encoding task, OL = object location task.

### Adjusting for Covariate Effects

In follow-up analyses, we entered age as a covariate into the correlational model. As before, in the brain-only model from the first set of analyses, age was negatively associated with all ROI factors except precuneus and with all method factors. Here, we regressed the ROI integrity factors, the method factors, and the episodic memory factor on age and examined the residual covariances between episodic memory and the ROI integrity factors (Figure 3). Age was negatively associated with episodic memory (Table 5). Moreover, the associations between episodic memory and the ROI factors were attenuated after controlling for age, leaving only hippocampus and parahippocampal gyrus being significantly associated with episodic memory, *r _EM, PFC_* = .17, Δ *χ^2^_(df = 1)_ = 2.69*; *p* = .10; *r _EM, HC_* = .29, Δ *χ^2^_(df = 1)_ = 11.30*; *p* = .0008; *r _EM, PHG_* = .28, Δ *χ^2^_(df = 1)_* = 8.10; *p* < .004; *r _EM, PRE_* = .07, Δ *χ^2^_(df = 1)_ = 0.62*; *p* = .43. Thus, the grey matter integrity factors of hippocampus and parahippocampal gyrus shared a significant amount of variance with episodic memory that was not collinear with age, further suggesting that the structural integrity of these two regions might be critical for episodic memory.

**Table 5.** Unique effects of age, education and sex on latent episodic memory and ROI factors

| Covariate | Effect on EM, $\beta$/<br>std. $\beta$ (SE) | Effect on PFC $\beta$/<br>std. $\beta$ (SE) | Effect on HC $\beta$/<br>std. $\beta$ (SE) | Effect on PHG<br>$\beta$/ std. $\beta$ (SE) | Effect on PRE $\beta$/ std.<br>$\beta$ (SE) |
| --- | --- | --- | --- | --- | --- |
| Age (years) | -0.06/-.22 (0.01)* | -0.02/-.15 (0.01) | -0.07/-.20 (0.02)* | -0.02/-.17 (0.01)* | -0.005/-.02 (0.02) |
| Education (years) | 0.08/.30 (0.01)* | 0.008/.08 (0.01) | 0.007/.02 (0.02) | -0.002/-.02 (0.01) | -0.03/-.13 (0.02) |
| Sex (men=0,<br>women=1) | .31* | .32* | .79* | .43* | .30* |

*Note. Standardized regression coefficient estimates and standard error (SE) of estimate. \*: $p < .05$ , according to standard errors/ Wald statistic. Outcome variables are standardized to mean = 5 and SD = 2. EM: episodic memory; PFC: prefrontal cortex; HC: hippocampus; PRE: precuneus. Sex differences are coded so that the effect quantifies female advantage if positive, and male advantage if negative.*

We then added years of education into the model as a covariate by regressing all ROI factors and episodic memory as well as the method factors on age and education, which had very little effect on estimates of associations and did not change which ones were significant, *r _EM, PFC_* = .17, Δ *χ^2^*_(df = 1)_ = 2.59; *p* = .11; *r _EM, HC_* = .29, Δ *χ^2^*_(df = 1)_ = 11.50; *p* < .0007; *r _EM, PHG_* = .29, Δ *χ^2^_(df = 1)_* = 8.88; *p* = .0028; *r _EM, PRE_* = .09, Δ *χ^2^_(df = 1)_* = 0.99; *p* = .32. When we added sex as a covariate, all associations were somewhat attenuated, but hippocampus and parahippocampal gyrus were still significantly associated with episodic memory, *r _EM, PFC_* = .08, Δ *χ^2^*_(df = 1)_ = 0.59; *p* = .44; *r _EM, HC_* = .21, Δ *χ^2^_(df = 1_*_)_ = 5.56; *p* = .018; *r _EM, PHG_* = .21, Δ *χ^2^_(df = 1)_* = 4.08; *p* = .043; *r _EM, PRE_* = .01, Δ *χ^2^_(df = 1)_* = 0.02; *p* = .88.

As an additional ad-hoc exploratory analysis, we investigated differences between men and women in the associations between grey matter integrity and episodic memory (supplementary material). After testing for measurement invariance across sexes, we ran the correlation model as a multigroup model, once with and once without age and education as covariates. The associations between hippocampal and parahippocampal grey matter integrity were restricted to men (see supplementary material for details).

## Discussion

In this study, we used cross-sectional data on multimodal structural imaging and episodic memory tasks from a large cohort to establish a structural equation model of regional grey matter structure integrity and its associations with episodic memory. We show that a MTMM latent factor representation of regional individual differences in grey matter structure enables researchers to examine links of structural brain properties to behavior. Specifically, this representation allows researchers to separate three sources of variance from one another:

i. variance shared within each ROI across imaging modalities (i.e., the ROI integrity factors);
ii. variance shared within each imaging modality across ROIs (i.e., the method factors); and
iii. variance unique to each ROI in each modality (i.e., residual variance).

The psychometric viability of the MTMM representation of regional grey matter integrity demonstrates that macro- and microstructural indicators of grey matter can indeed be combined to yield latent factors of grey matter integrity. In addition, the latent integrity factors formed a positive manifold, indicating that individual differences in grey matter integrity are correlated across regions. By moving away from specific aspects of integrity indicators to the expression of their common variance at the latent level, we pave the way for a deeper understanding of relations between brain structure and cognitive performance.

Age was negatively related to all ROI-wise latent integrity factors except precuneus, with older participants showing reduced grey matter integrity. This result is largely in line with previous findings based on single indicators focusing on volume (Raz et al. 2005; Fjell and Walhovd 2011), and MT (Seiler et al. 2014), or MD (Grydeland et al. 2013). Also, all method factors showed age differences. This suggests that the variance in each of the ROI factors is not capturing all age differences. There are still age differences in the methods’ unique variances. In other words, older individuals tend to have better grey matter integrity (across modalities) in prefrontal cortex, hippocampus and parahippocampal gyrus, and in addition, they tend to have smaller volumes, lower MT, and higher MD across ROIs.

To examine whether latent grey matter integrity factors are related to episodic memory, we tested their associations with episodic memory ability at the latent level. Episodic memory showed first-order associations with the structural integrity factors of all ROIs except precuneus, but not with the modality-specific method factors. When adjusted for age differences, hippocampus, and parahippocampal gyrus continued to be associated with episodic memory. That is, while prefrontal cortex’s first-order association with episodic memory could be accounted for by age differences in both grey matter structure and performance, the integrity of hippocampal and parahippocampal grey matter not only reflected individual differences collinear with chronological age, but also associations with episodic memory performance over and above age. Adjusting for inter-individual differences in years of education did not substantially affect the associations, with hippocampus and parahippocampal gyrus still showing the strongest associations with episodic memory. When adjusting for sex differences in episodic memory and ROI integrity, hippocampus and parahippocampal gyrus were still significantly associated with episodic memory. This is corroborated in the regression model, with hippocampus showing the numerically largest unique effect. Overall, this result strongly supports the hypothesis that maintained structural integrity of the hippocampus is germane to preserved episodic memory ability in old age (de Chastelaine et al. 2011; Nyberg et al. 2012; Cabeza et al. 2018; Nyberg and Lindenberger 2019; Nyberg and Pudas 2019).

The aims of this study were to establish a grey matter integrity factor model and validate it by associating its latent factors with episodic memory performance. We did not previously plan to investigate sex differences in measurement models or associations. Only after observing sex differences both in grey matter integrity and in episodic memory, we ran additional ad-hoc exploratory analyses to compare the models across men and women. We found the associations between hippocampal and parahippocampal grey matter integrity to be restricted to men. We provide details on these additional analyses and a short discussion in the supplementary material. In essence, we still generally hypothesize that the hippocampal and parahippocampal integrity in older adults are relevant for episodic memory performance irrespective of sex. At this point, we can only speculate that, there might be more men than women who have already experienced some grey matter integrity deterioration with consequences for memory functioning, possibly related to men carrying a higher metabolic risk with detrimental effects for both grey matter integrity and episodic memory (Raz et al., 2005; Raz and Rodrigue, 2006; Yates et al., 2012). This could also be a reason behind the observation that men show on average lower integrity in all ROIs and in episodic memory (Table 4). Further elaboration and investigation of these sex differences would exceed the scope of this study and should be pursued in future studies based on longitudinal data.

Our results also suggest that the combination of multimodal data yields information about general properties of grey matter tissue that are age-sensitive and relevant for older adults’ episodic memory performance, whereas little to none of this age- and memory-related information is captured by the specific variance of the methods themselves. This raises the important question which physiological aspects of grey matter are captured by the common variance of regional brain integrity as estimated by VBM, MT, and MD. Given that MD and MT load on the same factor as VBM, it seems worthwhile to consider physiological factors that affect the physical properties of the tissue and its overall size. Normal aging is marked by the loss of dendritic spines, dendritic arbors, synaptic density, and myelinated axons (Hof and Morrison 2004; Morrison and Baxter 2012); in addition, normal aging also involves loss of glia and small blood vessels (Raz and Daugherty 2018). All of these processes can be assumed to lead to a reduction in average tissue density as captured by MD and MT, and to a concomitant decrease in overall volume as captured by VBM. In terms of relative contributions to variations in the MR signal that affect MD, MT, and VBM in a correlated manner, we surmise that individual differences in cortical myelin might play a prominent role. Given that histochemical staining of myelin has shown that myelin coverage is more extensive in deeper relative to superficial cortical layers (Timmler and Simons 2019), one way to follow up on this proposition would be to test for differences in myelin content between layers using structural imaging methods with laminar resolution (Waehnert et al. 2014).

The MTMM model of regional grey matter integrity introduced in this article reflects correlated traits and correlated methods, and properly accounts for the non-independent structure of measurement errors in our data. Similar to other confirmatory factor analysis variants of structural equation modeling, it links the measurement model (ROI integrity factors, method factors, and residuals) to the structural model (associations between latent factors). The structural part of the model allows researchers to explore relations to behavior, and their modulation by covariates.

The proposed model is certainly not the only way to model associations between brain structure and cognition (Kievit et al. 2012). Hence, we would like to encourage researchers to adopt the MTMM approach whenever they have multiple measures for a given construct of interest. For example, models might capture a larger number of ROIs, a more fine-grained parcellation of ROIs into sub-regions, inter-hemispheric differences and commonalities, and a larger number of indicators. Furthermore, the general approach can be expanded to include factors of white-matter integrity, neurochemistry, or brain activity as assessed by functional MRI.

In this study, we have used a statistical approach to model the common variance across multiple indicators of grey matter integrity in latent factors for each ROI. At this point, we can only speculate about the physiological basis of individual differences in grey matter integrity captured with the MMMT approach. To overcome these ambiguities, the field needs a stronger coalition between animal models and human research, with structural MRI serving as a critical link (Lerch et al. 2017).

Furthermore, the present findings are based on cross-sectional data. The observed associations with age represent the joint outcome of individual differences in normal aging and more stable individual differences that were present in early adulthood (Hertzog 1985). It remains to be seen how individual differences in latent patterns of brain integrity change map onto changes in episodic memory. Hence, the present analyses need to be extended to longitudinal investigations that examine individual differences in latent brain integrity changes and their correlation with cognitive changes (for methodological work in developmental psychology, see Geiser et al. 2010).

Also note that this paper focuses on the association structure of individual differences in a healthy older population. Our results might not hold for all subgroups. Plausibly, there are hidden heterogeneities in the association structures that should be elucidated by follow-up studies. For instance, associations between dopamine availability and cognition have been found to differ between subgroups in a latent class analysis (Lövdén et al. 2017). Another data-driven way to identify hidden heterogeneities in associations are decision trees (Strobl et al. 2009), which can be usefully combined with structural equation models in structural equation modelling trees (Brandmaier et al. 2013). In addition to the structural integrity measures we investigated in this study, a comprehensive understanding of maintenance may further benefit from the integration of additional imaging modalities such as white matter integrity, neurochemical, and connectivity measures.

By applying MMMT modeling to data from a large sample of BASE-II participants, we established latent factors of grey matter integrity in hippocampus, parahippocampal gyrus, prefrontal cortex, and precuneus, which represented the shared variance of VBM, MT, and MD for each of these regions. Further, we found that older adults with greater structural integrity in hippocampus and parahippocampal gyrus also showed higher levels of episodic memory performance, with hippocampus contributing showing the largest unique association. Our results are consistent with the hypothesis that maintained structural integrity of the hippocampus helps to preserve episodic memory in old age. Future research needs to corroborate the content validity of the latent brain factors, and extend the present approach to longitudinal observations.

## Acknowledgement

We are grateful for the assistance of the MRI team at the Max Planck Berlin Institute for Human Development consisting of Sonali Beckmann, Nils Bodammer, Thomas Feg, Sebastian Schröder, and Nadine Taube, for the team leading the cognitive tests, and for all participants of BASE-II. We thank Julia Delius for copyediting.

## Data Availability Statement

Data can be requested from the steering committee of the Berlin Aging Study II. Further information regarding the application can be found under https://www.base2.mpg.de/en.

