## Supplemental Material for "Hippocampal and Parahippocampal Grey Matter Structural Integrity Assessed by Multimodal Imaging is Associated With Episodic Memory in Old Age"

[+]Joint last authorship

### 1. Episodic memory task descriptions

Scene Encoding. The scene encoding task was used to assess recognition of incidentally encoded scenes. Participants were presented with a series of grey-scale photos of indoor and outdoor scenes (500*300 pixels resolution, mean grey value 127. SD = 75. neutral valence). each presented for 2.5 s. Participants were instructed to decide for each scene whether it was indoor (button press with left index finger) or outdoor (right index finger). After 10 practice trials with each 5 indoor and outdoor scenes, participants were asked to view and judge 88 scenes (44 of each type). Then, they were presented with 44 of the seen items (22 indoor. 22 outdoor) randomly intermixed with 44 new items (22 indoor, 22 outdoor), and asked to indicate how confident they are that they had seen the scene before (on a scale from 1 = sure that new to 5 = sure that old). There was no time restriction on the response. After 2.5 hours, the recognition test was repeated (delayed recognition). In this study, we used the recognition score (hits minus false alarms) from the delayed recognition test.

Verbal Learning and Memory test (VLMT) The VLMT is a verbal list learning task that was used to assess recall of words after deliberate learning. Participants are presented a list of 15 words auditory via headphones for 5 times (always in the same order). After each presentation of the word list, participants are instructed to enter all words they can recall into the computer via keyboard. They enter the words one by one followed by enter, which lets the current word disappear so that only one word is visible at the time. After 5 times the word list, an interference list with 15 new words is presented and the participants are again requested to enter all words they can recall. Subsequently, participants are requested to enter all words they can recall from the first list, without the first list being presented again (free recall after distraction). After 30 minutes of performing other tasks, participants are asked to recall as many words as possible from the first list again (delayed recall), followed by a recognition test. In the recognition test, they were presented with all words from the first and the second list plus 20 new words, consisting of 5 of each of the following: 1) synonyms to words from the first and 2) from the second list, and 3) orthographically similar words to words from the first and 2) from the second list. The VLMT took about 30 minutes in total. After one week (same time of the day, seven days later), participants came back for further assessments, and were asked to recall and enter the words from the first list again, without the list being presented again. For the purpose of this study, we used the mean of recalled words across all 5 learning trials.

Face-Profession task The face profession task was used to assess associative memory recognition performance after incidental encoding. After 2 practice trials with encoding and recall. participants were requested to view 45 visually presented face-profession pairs and rate whether the face matches the profession. Next, they were given a distraction task in which they had to count down in intervals of three from 5679 for two minutes. After that, they were asked to watch a series of 54 face-profession pairs and to indicate for each pair whether they had seen the pair before. After their answer, they were asked to rate their confidence in the answer (not sure, quite sure, very sure). The set of 54 pairs in the recognition phase consisted of 27 old pairs, 9 new pairs, and 18 newly arranged pairs. For the purpose of this study, we used the recognition score (hits minus false alarms).

Object-Location task The object-location task was used to assess recall of deliberately encoded object locations. Sequences of 12 photographs of real-world objects were displayed at different locations in a 6 by 6 locations grid. After the sequence, objects appeared at the side of the screen and participants were asked to move pictures to the location they were first presented at, by means of clicking the picture and the location with the mouse. The task consisted of one practice trial and two test trials. The sum of correct placements across the two test trials is used in this study.

### 2. Parameter estimates of final models

Tables S1-S3 contain the parameter estimates, their standard errors of estimation, standardized estimates and their standard errors, z-values and p-values as in lavaan (Rosseel 2012) output. Table S1 contains estimates for the correlation model, Table S2 for the correlation model with covariates, and S3 for the regression model.

#### Table S1. Complete list of parameter estimates for the correlation model

| Latent variable | Loading | Observed variable | Estimate | SE | Estimate std. | SE std. | Z | P-value |
|---|---|---|---|---|---|---|---|---|
| PFC | =~ | Vmo | 1 | 0 | 0.3081 | 0.0667 | 4.6175 | <0.0001 |
| PFC | =~ | MTmo | 1.8753 | 0.4039 | 0.5728 | 0.0724 | 7.9124 | <0.0001 |
| PFC | =~ | MDmo | 0.1139 | 0.3543 | 0.0351 | 0.1052 | 0.3332 | 0.739 |
| PFC | =~ | Vdl | 0.4888 | 0.1998 | 0.1507 | 0.0764 | 1.973 | 0.0485 |
| PFC | =~ | MTdl | 0.3468 | 0.3002 | 0.1057 | 0.1002 | 1.0548 | 0.2915 |
| PFC | =~ | MDdl | -1.8911 | 0.4038 | -0.5714 | 0.088 | -6.4904 | <0.0001 |
| HC | =~ | Vhc | 1 | 0 | 0.6663 | 0.0345 | 19.3368 | <0.0001 |
| HC | =~ | MThc | 1.4262 | 0.0887 | 0.9416 | 0.0165 | 57.0327 | <0.0001 |
| HC | =~ | MDhc | -1.5291 | 0.0983 | -0.9205 | 0.0171 | -53.7624 | <0.0001 |
| PHG | =~ | Vphg | 1 | 0 | 0.3419 | 0.0564 | 6.0615 | <0.0001 |
| PHG | =~ | MTphg | 2.1727 | 0.3753 | 0.7215 | 0.045 | 16.0457 | <0.0001 |
| PHG | =~ | MDphg | -2.2859 | 0.3912 | -0.7398 | 0.0453 | -16.349 | <0.0001 |
| PRE | =~ | Vpre | 1 | 0 | 0.5604 | 0.0424 | 13.2259 | <0.0001 |
| PRE | =~ | MTpre | 1.3128 | 0.1262 | 0.7002 | 0.042 | 16.6695 | <0.0001 |
| PRE | =~ | MDpre | -1.1013 | 0.1116 | -0.6139 | 0.0433 | -14.1877 | <0.0001 |
| V | =~ | Vpre | 1 | 0 | 0.6385 | 0.0467 | 13.6768 | <0.0001 |
| V | =~ | Vphg | 0.6021 | 0.1072 | 0.3835 | 0.0578 | 6.6413 | <0.0001 |
| V | =~ | Vhc | 0.5636 | 0.0977 | 0.356 | 0.0521 | 6.8317 | <0.0001 |
| V | =~ | Vdl | 1.0951 | 0.1381 | 0.6926 | 0.0489 | 14.1682 | <0.0001 |
| V | =~ | Vmo | 0.8673 | 0.1269 | 0.5483 | 0.0572 | 9.5885 | <0.0001 |
| MT | =~ | MTpre | 1 | 0 | 0.6671 | 0.0398 | 16.7686 | <0.0001 |
| MT | =~ | MTphg | 0.6153 | 0.1004 | 0.4179 | 0.0629 | 6.6429 | <0.0001 |
| MT | =~ | MThc | 0.4378 | 0.074 | 0.3008 | 0.0484 | 6.2198 | <0.0001 |
| MT | =~ | MTdl | 1.2406 | 0.1128 | 0.8516 | 0.0333 | 25.5519 | <0.0001 |
| MT | =~ | MTmo | 0.8884 | 0.1203 | 0.6113 | 0.066 | 9.2585 | <0.0001 |
| MD | =~ | MDpre | 1 | 0 | 0.6938 | 0.0352 | 19.7207 | <0.0001 |
| MD | =~ | MDphg | 0.8096 | 0.0812 | 0.5334 | 0.0525 | 10.1595 | <0.0001 |
| MD | =~ | MDhc | 0.3764 | 0.0782 | 0.2347 | 0.0503 | 4.6637 | <0.0001 |
| MD | =~ | MDdl | 1.1082 | 0.1061 | 0.7506 | 0.0607 | 12.3593 | <0.0001 |
| MD | =~ | MDmo | 1.3557 | 0.1098 | 0.9351 | 0.0299 | 31.2265 | <0.0001 |
| EM | =~ | VLMT | 1 | 0 | 0.5252 | 0.0302 | 17.4134 | <0.0001 |

| | | | | | | | | |
|---|---|---|---|---|---|---|---|---|
| EM | =~ | FP | 0.8629 | 0.083 | 0.4532 | 0.0304 | 14.8967 | <0.0001 |
| EM | =~ | SE | 0.966 | 0.09 | 0.5074 | 0.0301 | 16.8324 | <0.0001 |
| EM | =~ | OL | 1.0835 | 0.098 | 0.5689 | 0.0303 | 18.7789 | <0.0001 |

| **Latent variable** | **Covariance** | **Observed variable** | **Estimate** | **SE** | **Estimate std.** | **SE std.** | **Z** | **P-value** |
|---|---|---|---|---|---|---|---|---|
| Vmo | ~~ | Vmo | 2.4139 | 0.25 | 0.6044 | 0.0588 | 10.2782 | <0.0001 |
| MTmo | ~~ | MTmo | 1.2119 | 0.2009 | 0.2982 | 0.0525 | 5.6821 | <0.0001 |
| MDmo | ~~ | MDmo | 0.4981 | 0.2274 | 0.1244 | 0.0576 | 2.1577 | 0.031 |
| Vdl | ~~ | Vdl | 1.9856 | 0.2541 | 0.4976 | 0.0637 | 7.813 | <0.0001 |
| MTdl | ~~ | MTdl | 1.0764 | 0.1974 | 0.2636 | 0.0515 | 5.1172 | <0.0001 |
| MDdl | ~~ | MDdl | 0.457 | 0.159 | 0.1101 | 0.04 | 2.7498 | 0.006 |
| Vhc | ~~ | Vhc | 1.7174 | 0.1609 | 0.4293 | 0.0409 | 10.5079 | <0.0001 |
| MThc | ~~ | MThc | 0.0936 | 0.0687 | 0.023 | 0.017 | 1.3491 | 0.1773 |
| MDhc | ~~ | MDhc | 0.4783 | 0.0943 | 0.0976 | 0.0209 | 4.6703 | <0.0001 |
| Vphg | ~~ | Vphg | 2.8956 | 0.2499 | 0.736 | 0.0482 | 15.2571 | <0.0001 |
| MTphg | ~~ | MTphg | 1.2713 | 0.1813 | 0.3049 | 0.0472 | 6.4617 | <0.0001 |
| MDphg | ~~ | MDphg | 0.7379 | 0.1565 | 0.1681 | 0.0386 | 4.3505 | <0.0001 |
| Vpre | ~~ | Vpre | 1.0896 | 0.1919 | 0.2783 | 0.0516 | 5.3938 | <0.0001 |
| MTpre | ~~ | MTpre | 0.2797 | 0.1879 | 0.0647 | 0.0447 | 1.4461 | 0.1482 |
| MDpre | ~~ | MDpre | 0.5612 | 0.1281 | 0.1418 | 0.0334 | 4.2394 | <0.0001 |
| VLMT | ~~ | VLMT | 2.8932 | 0.1452 | 0.7241 | 0.0317 | 22.8526 | <0.0001 |
| FP | ~~ | FP | 3.1758 | 0.1418 | 0.7946 | 0.0276 | 28.8177 | <0.0001 |
| SE | ~~ | SE | 2.967 | 0.1438 | 0.7426 | 0.0306 | 24.2748 | <0.0001 |
| OL | ~~ | OL | 2.7044 | 0.1497 | 0.6764 | 0.0345 | 19.6222 | <0.0001 |

| **Latent variable** | **Covariance** | **Latent variable** | **Estimate** | **SE** | **Estimate std.** | **SE std.** | **Z** | **P-value** |
|---|---|---|---|---|---|---|---|---|
| PFC | ~~ | PFC | 0.3791 | 0.171 | 1 | 0 | NA | NA |
| HC | ~~ | HC | 1.7761 | 0.2627 | 1 | 0 | NA | NA |
| PHG | ~~ | PHG | 0.4598 | 0.1601 | 1 | 0 | NA | NA |
| PRE | ~~ | PRE | 1.2298 | 0.2156 | 1 | 0 | NA | NA |
| V | ~~ | V | 1.5965 | 0.2821 | 1 | 0 | NA | NA |
| MD | ~~ | MD | 1.9055 | 0.2843 | 1 | 0 | NA | NA |
| MT | ~~ | MT | 1.9239 | 0.3263 | 1 | 0 | NA | NA |
| EM | ~~ | EM | 1.1023 | 0.1414 | 1 | 0 | NA | NA |

| **Latent variable** | **Covariance** | **Latent variable** | **Estimate** | **SE** | **Estimate std.** | **SE std.** | **Z** | **P-value** |
|---|---|---|---|---|---|---|---|---|
| PFC | ~~ | HC | 0.4817 | 0.1468 | 0.587 | 0.061 | 9.6168 | <0.0001 |
| PFC | ~~ | PHG | 0.2594 | 0.1002 | 0.6213 | 0.0751 | 8.269 | <0.0001 |
| PFC | ~~ | PRE | -0.0538 | 0.0844 | -0.0788 | 0.1303 | -0.6049 | 0.5452 |
| HC | ~~ | PHG | 0.7428 | 0.178 | 0.822 | 0.0332 | 24.7335 | <0.0001 |
| HC | ~~ | PRE | 0.0353 | 0.1279 | 0.0239 | 0.0861 | 0.2772 | 0.7817 |
| PHG | ~~ | PRE | 0.1812 | 0.0806 | 0.241 | 0.087 | 2.7711 | 0.0056 |
| V | ~~ | MT | 0.8186 | 0.1964 | 0.4671 | 0.0761 | 6.1383 | <0.0001 |
| V | ~~ | MD | -1.1448 | 0.2067 | -0.6564 | 0.0518 | -12.6669 | <0.0001 |
| MT | ~~ | MD | -1.5627 | 0.2516 | -0.8162 | 0.0342 | -23.8638 | <0.0001 |
| PFC | ~~ | EM | 0.1575 | 0.0753 | 0.2436 | 0.0976 | 2.4967 | 0.0125 |

| | | | | | | | | |
|---|---|---|---|---|---|---|---|---|
| HC | ~~ | EM | 0.4759 | 0.1206 | 0.3401 | 0.0764 | 4.453 | <0.0001 |
| PHG | ~~ | EM | 0.234 | 0.076 | 0.3287 | 0.0878 | 3.7453 | 0.0002 |
| PRE | ~~ | EM | 0.1124 | 0.1103 | 0.0965 | 0.0944 | 1.0225 | 0.3065 |
| **Observed variable** | **Residual covariance** | **Observed variable** | **Estimate** | **SE** | **Estimate std.** | **SE std.** | **Z** | **P-value** |
| Vmo | ~~ | Vdl | 0.6093 | 0.1987 | 0.2783 | 0.0723 | 3.849 | 0.0001 |
| Vhc | ~~ | Vphg | 1.4039 | 0.1707 | 0.6296 | 0.0389 | 16.1732 | <0.0001 |
| **Observed variable** | **Intercept** | | **Estimate** | **SE** | **Estimate std.** | **SE std.** | **Z** | **P-value** |
| Vmo | ~1 | | 4.9921 | 0.1099 | 2.498 | 0.1118 | 22.3525 | <0.0001 |
| MTmo | ~1 | | 4.9343 | 0.1298 | 2.4477 | 0.1371 | 17.8512 | <0.0001 |
| MDmo | ~1 | | 5.0246 | 0.1175 | 2.5107 | 0.1211 | 20.7388 | <0.0001 |
| Vdl | ~1 | | 4.9962 | 0.11 | 2.501 | 0.1119 | 22.3541 | <0.0001 |
| MTdl | ~1 | | 4.948 | 0.132 | 2.4487 | 0.1391 | 17.6049 | <0.0001 |
| MDdl | ~1 | | 5.1117 | 0.1183 | 2.5084 | 0.1179 | 21.2707 | <0.0001 |
| Vhc | ~1 | | 4.9763 | 0.109 | 2.4879 | 0.1091 | 22.7988 | <0.0001 |
| MThc | ~1 | | 4.9551 | 0.1158 | 2.4546 | 0.1222 | 20.0862 | <0.0001 |
| MDhc | ~1 | | 5.2121 | 0.1234 | 2.3543 | 0.1116 | 21.0994 | <0.0001 |
| Vphg | ~1 | | 4.9883 | 0.1089 | 2.515 | 0.1108 | 22.697 | <0.0001 |
| MTphg | ~1 | | 4.9037 | 0.1292 | 2.4013 | 0.1348 | 17.8079 | <0.0001 |
| MDphg | ~1 | | 5.167 | 0.1202 | 2.4662 | 0.1156 | 21.3371 | <0.0001 |
| Vpre | ~1 | | 4.9944 | 0.109 | 2.5239 | 0.1091 | 23.144 | <0.0001 |
| MTpre | ~1 | | 4.96 | 0.1285 | 2.3855 | 0.1256 | 18.9952 | <0.0001 |
| MDpre | ~1 | | 5.0412 | 0.1145 | 2.5338 | 0.1166 | 21.7249 | <0.0001 |
| VLMT | ~1 | | 5.0015 | 0.0516 | 2.5022 | 0.0525 | 47.6818 | <0.0001 |
| FP | ~1 | | 5.0003 | 0.0517 | 2.5012 | 0.0525 | 47.6183 | <0.0001 |
| SE | ~1 | | 5.0015 | 0.0517 | 2.5021 | 0.0525 | 47.6474 | <0.0001 |
| OL | ~1 | | 4.9994 | 0.0515 | 2.5001 | 0.0524 | 47.7513 | <0.0001 |
| **Latent variable** | **Mean** | | **Estimate** | **SE** | **Estimate std.** | **SE std.** | **Z** | **P-value** |
| PFC | ~1 | | 0 | 0 | 0 | 0 | NA | NA |
| HC | ~1 | | 0 | 0 | 0 | 0 | NA | NA |
| PHG | ~1 | | 0 | 0 | 0 | 0 | NA | NA |
| PRE | ~1 | | 0 | 0 | 0 | 0 | NA | NA |
| V | ~1 | | 0 | 0 | 0 | 0 | NA | NA |
| MT | ~1 | | 0 | 0 | 0 | 0 | NA | NA |
| MD | ~1 | | 0 | 0 | 0 | 0 | NA | NA |
| EM | ~1 | | 0 | 0 | 0 | 0 | NA | NA |

*Note. P-values from Wald test.*

Table S2. Complete list of parameter estimates for the correlation model with age, education, and sex as covariates

| Latent variable | Loading | Observed variable | Estimate | SE | Estimate std. | SE std. | Z | P-value |
|---|---|---|---|---|---|---|---|---|
| PFC | =~ | Vmo | 1 | 0 | 0.185 | 0.0719 | 2.5716 | 0.0101 |
| PFC | =~ | MTmo | 2.288 | 0.814 | 0.4203 | 0.0884 | 4.7519 | <0.0001 |
| PFC | =~ | MDmo | 1.445 | 0.9764 | 0.2688 | 0.105 | 2.561 | 0.0104 |
| PFC | =~ | Vdl | 0.0477 | 0.4223 | 0.0088 | 0.0804 | 0.1098 | 0.9126 |
| PFC | =~ | MTdl | -0.5225 | 0.7032 | -0.0955 | 0.105 | -0.9089 | 0.3634 |
| PFC | =~ | MDdl | -2.1415 | 0.7377 | -0.3914 | 0.1004 | -3.8968 | 0.0001 |
| HC | =~ | Vhc | 1 | 0 | 0.6083 | 0.0376 | 16.1651 | <0.0001 |
| HC | =~ | MThc | 1.472 | 0.0993 | 0.8921 | 0.024 | 37.1789 | <0.0001 |
| HC | =~ | MDhc | -1.5964 | 0.111 | -0.881 | 0.0235 | -37.5033 | <0.0001 |
| PHG | =~ | Vphg | 1 | 0 | 0.2733 | 0.0582 | 4.6983 | <0.0001 |
| PHG | =~ | MTphg | 2.3987 | 0.5191 | 0.6389 | 0.0532 | 12.0154 | <0.0001 |
| PHG | =~ | MDphg | -2.6318 | 0.5769 | -0.6808 | 0.0467 | -14.5803 | <0.0001 |
| PRE | =~ | Vpre | 1 | 0 | 0.5597 | 0.0414 | 13.5317 | <0.0001 |
| PRE | =~ | MTpre | 1.3375 | 0.1254 | 0.7144 | 0.0395 | 18.0779 | <0.0001 |
| PRE | =~ | MDpre | -1.1527 | 0.1129 | -0.6437 | 0.0428 | -15.0379 | <0.0001 |
| V | =~ | Vpre | 1 | 0 | 0.5883 | 0.0479 | 12.2892 | <0.0001 |
| V | =~ | Vphg | 0.6645 | 0.1192 | 0.3917 | 0.0576 | 6.8056 | <0.0001 |
| V | =~ | Vhc | 0.6541 | 0.1141 | 0.3822 | 0.0529 | 7.2239 | <0.0001 |
| V | =~ | Vdl | 1.2197 | 0.1486 | 0.7152 | 0.0457 | 15.6563 | <0.0001 |
| V | =~ | Vmo | 0.9754 | 0.1395 | 0.5713 | 0.0563 | 10.1405 | <0.0001 |
| MT | =~ | MTpre | 1 | 0 | 0.6256 | 0.0451 | 13.8772 | <0.0001 |
| MT | =~ | MTphg | 0.6833 | 0.1168 | 0.4374 | 0.0643 | 6.8007 | <0.0001 |
| MT | =~ | MThc | 0.4814 | 0.0868 | 0.3123 | 0.0499 | 6.2569 | <0.0001 |
| MT | =~ | MTdl | 1.3282 | 0.1281 | 0.8561 | 0.0321 | 26.6309 | <0.0001 |
| MT | =~ | MTmo | 1.0497 | 0.1512 | 0.6802 | 0.0695 | 9.7897 | <0.0001 |
| MD | =~ | MDpre | 1 | 0 | 0.6538 | 0.0395 | 16.5333 | <0.0001 |
| MD | =~ | MDphg | 0.845 | 0.0942 | 0.525 | 0.0515 | 10.1884 | <0.0001 |
| MD | =~ | MDhc | 0.4146 | 0.0898 | 0.2448 | 0.0517 | 4.7379 | <0.0001 |
| MD | =~ | MDdl | 1.2586 | 0.1339 | 0.811 | 0.0635 | 12.775 | <0.0001 |
| MD | =~ | MDmo | 1.4403 | 0.1187 | 0.9449 | 0.0379 | 24.9273 | <0.0001 |
| EM | =~ | VLMT | 1 | 0 | 0.5501 | 0.0289 | 19.049 | <0.0001 |
| EM | =~ | FP | 0.8633 | 0.079 | 0.475 | 0.0294 | 16.1681 | <0.0001 |
| EM | =~ | SE | 0.901 | 0.0809 | 0.4957 | 0.0291 | 17.0245 | <0.0001 |
| EM | =~ | OL | 0.9754 | 0.0839 | 0.5364 | 0.0291 | 18.4611 | <0.0001 |
| **Observed variable** | **Residual variance** | | **Estimate** | **SE** | **Estimate std.** | **SE std.** | **Z** | **P-value** |
| Vmo | ~~ | Vmo | 2.4368 | 0.2521 | 0.6072 | 0.059 | 10.285 | <0.0001 |
| MTmo | ~~ | MTmo | 1.1512 | 0.198 | 0.2829 | 0.0517 | 5.4758 | <0.0001 |
| MDmo | ~~ | MDmo | 0.4038 | 0.2373 | 0.1018 | 0.0603 | 1.6869 | 0.0916 |
| Vdl | ~~ | Vdl | 1.9479 | 0.2456 | 0.4865 | 0.0616 | 7.8928 | <0.0001 |
| MTdl | ~~ | MTdl | 1.1522 | 0.2033 | 0.2802 | 0.0529 | 5.2914 | <0.0001 |

| | | | | | | | | |
|---|---|---|---|---|---|---|---|---|
| MDdl | ~~ | MDdl | 0.4344 | 0.1594 | 0.1057 | 0.0401 | 2.6325 | 0.0085 |
| Vhc | ~~ | Vhc | 1.6474 | 0.1584 | 0.4086 | 0.0408 | 10.0174 | <0.0001 |
| MThc | ~~ | MThc | 0.0963 | 0.0699 | 0.0237 | 0.0174 | 1.3637 | 0.1727 |
| MDhc | ~~ | MDhc | 0.474 | 0.0967 | 0.0968 | 0.0213 | 4.5438 | <0.0001 |
| Vphg | ~~ | Vphg | 2.91 | 0.2488 | 0.7344 | 0.0483 | 15.1922 | <0.0001 |
| MTphg | ~~ | MTphg | 1.3226 | 0.1861 | 0.3171 | 0.048 | 6.6045 | <0.0001 |
| MDphg | ~~ | MDphg | 0.6709 | 0.1727 | 0.1517 | 0.0414 | 3.6652 | 0.0002 |
| Vpre | ~~ | Vpre | 1.2288 | 0.1818 | 0.309 | 0.0482 | 6.4083 | <0.0001 |
| MTpre | ~~ | MTpre | 0.3441 | 0.1933 | 0.0788 | 0.0457 | 1.7229 | 0.0849 |
| MDpre | ~~ | MDpre | 0.5589 | 0.1321 | 0.1399 | 0.0344 | 4.0686 | <0.0001 |
| VLMT | ~~ | VLMT | 2.7895 | 0.142 | 0.6973 | 0.0318 | 21.9455 | <0.0001 |
| FP | ~~ | FP | 3.0979 | 0.1396 | 0.7744 | 0.0279 | 27.7527 | <0.0001 |
| SE | ~~ | SE | 3.017 | 0.1397 | 0.7543 | 0.0289 | 26.1289 | <0.0001 |
| OL | ~~ | OL | 2.8518 | 0.1416 | 0.7123 | 0.0312 | 22.8557 | <0.0001 |
| **Latent variable** | **Variance** | | **Estimate** | **SE** | **Estimate std.** | **SE std.** | **Z** | **P-value** |
| PFC | ~~ | PFC | 0.1232 | 0.0953 | 0.8974 | 0.0519 | 17.2812 | <0.0001 |
| HC | ~~ | HC | 1.3211 | 0.2013 | 0.8857 | 0.0384 | 23.0741 | <0.0001 |
| PHG | ~~ | PHG | 0.2572 | 0.1129 | 0.8692 | 0.0479 | 18.1411 | <0.0001 |
| PRE | ~~ | PRE | 1.2099 | 0.2137 | 0.9713 | 0.0206 | 47.0837 | <0.0001 |
| V | ~~ | V | 1.049 | 0.216 | 0.762 | 0.0596 | 12.7834 | <0.0001 |
| MD | ~~ | MD | 1.2564 | 0.2039 | 0.7359 | 0.0474 | 15.5187 | <0.0001 |
| MT | ~~ | MT | 1.3589 | 0.2562 | 0.7952 | 0.0564 | 14.0965 | <0.0001 |
| EM | ~~ | EM | 1.0225 | 0.1235 | 0.8446 | 0.0249 | 33.8634 | <0.0001 |
| **Covariate** | **Variance** | | **Estimate** | **SE** | **Estimate std.** | **SE std.** | **Z** | **P-value** |
| age | ~~ | age | 14.7178 | 0.5353 | 1 | 0 | NA | NA |
| edu | ~~ | edu | 17.3996 | 0.6724 | 1 | 0 | NA | NA |
| sex | ~~ | sex | 0.2499 | 0.0091 | 1 | 0 | NA | NA |
| **Latent variable** | **Residual covariance** | **Latent variable** | **Estimate** | **SE** | **Estimate std.** | **SE std.** | **Z** | **P-value** |
| PFC | ~~ | HC | 0.1871 | 0.0889 | 0.4638 | 0.0718 | 6.4627 | <0.0001 |
| PFC | ~~ | PHG | 0.0756 | 0.0482 | 0.4245 | 0.1068 | 3.974 | 0.0001 |
| PFC | ~~ | PRE | -0.1284 | 0.052 | -0.3325 | 0.1149 | -2.8926 | 0.0038 |
| HC | ~~ | PHG | 0.442 | 0.1276 | 0.7583 | 0.0453 | 16.7536 | <0.0001 |
| HC | ~~ | PRE | -0.1703 | 0.0926 | -0.1347 | 0.0715 | -1.8831 | 0.0597 |
| PHG | ~~ | PRE | 0.0726 | 0.0473 | 0.1302 | 0.0772 | 1.6858 | 0.0918 |
| PFC | ~~ | EM | 0.4335 | 0.1304 | 0.3631 | 0.0852 | 4.2604 | <0.0001 |
| HC | ~~ | EM | -0.7036 | 0.1418 | -0.6129 | 0.0582 | -10.5285 | <0.0001 |
| PHG | ~~ | EM | -1.0187 | 0.1781 | -0.7796 | 0.0394 | -19.7698 | <0.0001 |
| PRE | ~~ | EM | 0.1871 | 0.0889 | 0.4638 | 0.0718 | 6.4627 | <0.0001 |
| V | ~~ | MT | 0.0756 | 0.0482 | 0.4245 | 0.1068 | 3.974 | 0.0001 |
| V | ~~ | MD | -0.1284 | 0.052 | -0.3325 | 0.1149 | -2.8926 | 0.0038 |
| MT | ~~ | MD | 0.442 | 0.1276 | 0.7583 | 0.0453 | 16.7536 | <0.0001 |
| **Observed variable** | **Residual covariance** | **Observed variable** | **Estimate** | **SE** | **Estimate std.** | **SE std.** | **Z** | **P-value** |
| Vmo | ~~ | Vdl | 0.6002 | 0.1975 | 0.2755 | 0.0726 | 3.7938 | 0.0001 |

| | | | | | | | | |
|---|---|---|---|---|---|---|---|---|
| Vhc | ~~ | Vphg | 1.3693 | 0.1687 | 0.6254 | 0.0393 | 15.9107 | <0.0001 |
| **Latent variable** | **Regression** | **Covariate** | **Estimate** | **SE** | **Estimate std.** | **SE std.** | **Z** | **P-value** |
| EM | ~ | age | -0.0628 | 0.0098 | -0.2188 | 0.032 | -6.8322 | <0.0001 |
| EM | ~ | sex | 0.3151 | 0.0741 | 0.1432 | 0.032 | 4.48 | <0.0001 |
| EM | ~ | edu | 0.0778 | 0.01 | 0.295 | 0.0332 | 8.8763 | <0.0001 |
| PFC | ~ | age | -0.0146 | 0.0114 | -0.1507 | 0.0859 | -1.7544 | 0.0794 |
| PFC | ~ | sex | 0.2 | 0.1032 | 0.2698 | 0.072 | 3.7446 | 0.0002 |
| PFC | ~ | edu | 0.0075 | 0.0071 | 0.0842 | 0.0729 | 1.1553 | 0.248 |
| HC | ~ | age | -0.0642 | 0.0194 | -0.2017 | 0.0564 | -3.5761 | 0.0003 |
| HC | ~ | sex | 0.6604 | 0.157 | 0.2704 | 0.0568 | 4.7624 | <0.0001 |
| HC | ~ | edu | 0.0068 | 0.0168 | 0.0232 | 0.0574 | 0.4041 | 0.6861 |
| PHG | ~ | age | -0.0241 | 0.0116 | -0.1697 | 0.0692 | -2.4509 | 0.0143 |
| PHG | ~ | sex | 0.347 | 0.1035 | 0.319 | 0.0645 | 4.9462 | <0.0001 |
| PHG | ~ | edu | -0.0021 | 0.0086 | -0.016 | 0.0657 | -0.244 | 0.8072 |
| PRE | ~ | age | 0.0054 | 0.0195 | 0.0186 | 0.0669 | 0.2782 | 0.7809 |
| PRE | ~ | sex | 0.2787 | 0.1585 | 0.1248 | 0.0708 | 1.764 | 0.0777 |
| PRE | ~ | edu | -0.0302 | 0.0179 | -0.1128 | 0.0661 | -1.7055 | 0.0881 |
| V | ~ | age | -0.0645 | 0.0196 | -0.2108 | 0.0614 | -3.4311 | 0.0006 |
| V | ~ | sex | 1.0306 | 0.1692 | 0.4392 | 0.0613 | 7.1628 | <0.0001 |
| V | ~ | edu | 0.0071 | 0.0187 | 0.0253 | 0.0662 | 0.3824 | 0.7021 |
| MT | ~ | age | -0.1149 | 0.0242 | -0.3371 | 0.061 | -5.5216 | <0.0001 |
| MT | ~ | sex | 0.7639 | 0.1927 | 0.2922 | 0.0668 | 4.3737 | <0.0001 |
| MT | ~ | edu | 0.0238 | 0.0215 | 0.0761 | 0.068 | 1.1184 | 0.2634 |
| MD | ~ | age | 0.1556 | 0.0207 | 0.4569 | 0.0462 | 9.8933 | <0.0001 |
| MD | ~ | sex | -0.6143 | 0.1603 | -0.235 | 0.0579 | -4.0602 | <0.0001 |
| MD | ~ | edu | 0.0024 | 0.0181 | 0.0078 | 0.0578 | 0.1344 | 0.8931 |
| **Observed variable** | **intercept** | | **Estimate** | **SE** | **Estimate std.** | **SE std.** | **Z** | **P-value** |
| Vmo | ~1 | | 5.1183 | 0.1078 | 2.555 | 0.1091 | 23.4234 | <0.0001 |
| MTmo | ~1 | | 5.0428 | 0.1261 | 2.4997 | 0.1334 | 18.7443 | <0.0001 |
| MDmo | ~1 | | 5.0106 | 0.1116 | 2.5157 | 0.1178 | 21.3487 | <0.0001 |
| Vdl | ~1 | | 5.1265 | 0.1079 | 2.562 | 0.109 | 23.5106 | <0.0001 |
| MTdl | ~1 | | 5.0086 | 0.1314 | 2.4697 | 0.1368 | 18.0469 | <0.0001 |
| MDdl | ~1 | | 5.0293 | 0.1081 | 2.4802 | 0.1148 | 21.5963 | <0.0001 |
| Vhc | ~1 | | 5.1098 | 0.1066 | 2.5449 | 0.1056 | 24.107 | <0.0001 |
| MThc | ~1 | | 5.0755 | 0.113 | 2.5186 | 0.1173 | 21.4646 | <0.0001 |
| MDhc | ~1 | | 5.095 | 0.121 | 2.3023 | 0.1108 | 20.7814 | <0.0001 |
| Vphg | ~1 | | 5.0956 | 0.1077 | 2.5598 | 0.109 | 23.4787 | <0.0001 |
| MTphg | ~1 | | 5.0269 | 0.1263 | 2.4614 | 0.1315 | 18.7236 | <0.0001 |
| MDphg | ~1 | | 5.0498 | 0.1132 | 2.4015 | 0.1137 | 21.1247 | <0.0001 |
| Vpre | ~1 | | 5.137 | 0.1082 | 2.576 | 0.1067 | 24.143 | <0.0001 |
| MTpre | ~1 | | 5.0617 | 0.1286 | 2.4225 | 0.1241 | 19.5145 | <0.0001 |
| MDpre | ~1 | | 4.9705 | 0.1122 | 2.4871 | 0.1158 | 21.4846 | <0.0001 |
| VLMT | ~1 | | 5.0018 | 0.0516 | 2.5009 | 0.0525 | 47.6483 | <0.0001 |

| | | | | | | | | |
|---|---|---|---|---|---|---|---|---|
| FP | ~1 | | 5.0005 | 0.0517 | 2.5001 | 0.0525 | 47.5902 | <0.0001 |
| SE | ~1 | | 5.0015 | 0.0517 | 2.5008 | 0.0525 | 47.6122 | <0.0001 |
| OL | ~1 | | 4.9992 | 0.0515 | 2.4985 | 0.0524 | 47.7062 | <0.0001 |
| age_c | ~1 | | -0.0022 | 0.0987 | -0.0006 | 0.0257 | -0.0219 | 0.9826 |
| edu_c | ~1 | | -0.0045 | 0.1136 | -0.0011 | 0.0272 | -0.04 | 0.9681 |
| **Latent variable** | **intercept** | | **Estimate** | **SE** | **Estimate std.** | **SE std.** | **Z** | **P-value** |
| PFC | ~1 | | 0 | 0 | 0 | 0 | NA | NA |
| HC | ~1 | | 0 | 0 | 0 | 0 | NA | NA |
| PHG | ~1 | | 0 | 0 | 0 | 0 | NA | NA |
| PRE | ~1 | | 0 | 0 | 0 | 0 | NA | NA |
| V | ~1 | | 0 | 0 | 0 | 0 | NA | NA |
| MT | ~1 | | 0 | 0 | 0 | 0 | NA | NA |
| MD | ~1 | | 0 | 0 | 0 | 0 | NA | NA |
| EM | ~1 | | 0 | 0 | 0 | 0 | NA | NA |
| age | ~1 | | 0 | 0 | 0 | 0 | NA | NA |
| edu | ~1 | | 0 | 0 | 0 | 0 | NA | NA |

*Note. P-values from Wald test.*

### Table S3. Complete list of parameter estimates for the regression model

| Latent variable | Loading | Observed variable | Estimate | SE | Estimate std. | SE std. | Z | P-value |
|---|---|---|---|---|---|---|---|---|
| PFC | =~ | Vmo | 1 | 0 | 0.3081 | 0.0667 | 4.6175 | <0.0001 |
| PFC | =~ | MTmo | 1.8754 | 0.404 | 0.5728 | 0.0724 | 7.9126 | <0.0001 |
| PFC | =~ | MDmo | 0.1139 | 0.3543 | 0.035 | 0.1052 | 0.3332 | 0.739 |
| PFC | =~ | Vdl | 0.4888 | 0.1998 | 0.1507 | 0.0764 | 1.9731 | 0.0485 |
| PFC | =~ | MTdl | 0.3468 | 0.3002 | 0.1057 | 0.1002 | 1.0548 | 0.2915 |
| PFC | =~ | MDdl | -1.8911 | 0.4038 | -0.5714 | 0.088 | -6.4905 | <0.0001 |
| HC | =~ | Vhc | 1 | 0 | 0.6663 | 0.0345 | 19.337 | <0.0001 |
| HC | =~ | MThc | 1.4262 | 0.0887 | 0.9416 | 0.0165 | 57.0333 | <0.0001 |
| HC | =~ | MDhc | -1.5291 | 0.0983 | -0.9205 | 0.0171 | -53.7627 | <0.0001 |
| PHG | =~ | Vphg | 1 | 0 | 0.3419 | 0.0564 | 6.0616 | <0.0001 |
| PHG | =~ | MTphg | 2.1727 | 0.3753 | 0.7215 | 0.045 | 16.0459 | <0.0001 |
| PHG | =~ | MDphg | -2.2858 | 0.3912 | -0.7398 | 0.0453 | -16.3491 | <0.0001 |
| PRE | =~ | Vpre | 1 | 0 | 0.5604 | 0.0424 | 13.2258 | <0.0001 |
| PRE | =~ | MTpre | 1.3128 | 0.1262 | 0.7002 | 0.042 | 16.6695 | <0.0001 |
| PRE | =~ | MDpre | -1.1013 | 0.1116 | -0.6139 | 0.0433 | -14.1877 | <0.0001 |
| V | =~ | Vpre | 1 | 0 | 0.6385 | 0.0467 | 13.6768 | <0.0001 |
| V | =~ | Vphg | 0.6021 | 0.1072 | 0.3835 | 0.0578 | 6.6413 | <0.0001 |
| V | =~ | Vhc | 0.5636 | 0.0977 | 0.356 | 0.0521 | 6.8318 | <0.0001 |
| V | =~ | Vdl | 1.0951 | 0.1381 | 0.6926 | 0.0489 | 14.1683 | <0.0001 |
| V | =~ | Vmo | 0.8673 | 0.1269 | 0.5483 | 0.0572 | 9.5885 | <0.0001 |
| MT | =~ | MTpre | 1 | 0 | 0.6671 | 0.0398 | 16.7685 | <0.0001 |
| MT | =~ | MTphg | 0.6153 | 0.1004 | 0.4179 | 0.0629 | 6.6429 | <0.0001 |

| | | | | | | | | |
|---|---|---|---|---|---|---|---|---|
| MT | =~ | MThc | 0.4378 | 0.074 | 0.3008 | 0.0484 | 6.2199 | <0.0001 |
| MT | =~ | MTdl | 1.2406 | 0.1128 | 0.8516 | 0.0333 | 25.5519 | <0.0001 |
| MT | =~ | MTmo | 0.8885 | 0.1203 | 0.6113 | 0.066 | 9.2586 | <0.0001 |
| MD | =~ | MDpre | 1 | 0 | 0.6938 | 0.0352 | 19.7205 | <0.0001 |
| MD | =~ | MDphg | 0.8096 | 0.0812 | 0.5334 | 0.0525 | 10.1595 | <0.0001 |
| MD | =~ | MDhc | 0.3764 | 0.0782 | 0.2347 | 0.0503 | 4.6637 | <0.0001 |
| MD | =~ | MDdl | 1.1082 | 0.1061 | 0.7506 | 0.0607 | 12.3593 | <0.0001 |
| MD | =~ | MDmo | 1.3557 | 0.1098 | 0.9351 | 0.0299 | 31.2265 | <0.0001 |
| EM | =~ | VLMT | 1 | 0 | 0.5252 | 0.0302 | 17.4136 | <0.0001 |
| EM | =~ | FP | 0.8629 | 0.083 | 0.4532 | 0.0304 | 14.8967 | <0.0001 |
| EM | =~ | SE | 0.966 | 0.09 | 0.5074 | 0.0301 | 16.8323 | <0.0001 |
| EM | =~ | OL | 1.0835 | 0.098 | 0.5689 | 0.0303 | 18.779 | <0.0001 |
| **Observed variable** | **Residual variance** | | **Estimate** | **SE** | **Estimate std.** | **SE std.** | **Z** | **P-value** |
| Vmo | ~~ | Vmo | 2.4139 | 0.25 | 0.6044 | 0.0588 | 10.2781 | <0.0001 |
| MTmo | ~~ | MTmo | 1.2119 | 0.2009 | 0.2982 | 0.0525 | 5.6821 | <0.0001 |
| MDmo | ~~ | MDmo | 0.4981 | 0.2274 | 0.1244 | 0.0576 | 2.1577 | 0.031 |
| Vdl | ~~ | Vdl | 1.9856 | 0.2541 | 0.4976 | 0.0637 | 7.813 | <0.0001 |
| MTdl | ~~ | MTdl | 1.0764 | 0.1974 | 0.2636 | 0.0515 | 5.1171 | <0.0001 |
| MDdl | ~~ | MDdl | 0.457 | 0.159 | 0.11 | 0.04 | 2.7497 | 0.006 |
| Vhc | ~~ | Vhc | 1.7174 | 0.1609 | 0.4293 | 0.0409 | 10.5078 | <0.0001 |
| MThc | ~~ | MThc | 0.0936 | 0.0687 | 0.023 | 0.017 | 1.3491 | 0.1773 |
| MDhc | ~~ | MDhc | 0.4783 | 0.0943 | 0.0976 | 0.0209 | 4.6704 | <0.0001 |
| Vphg | ~~ | Vphg | 2.8956 | 0.2499 | 0.736 | 0.0482 | 15.257 | <0.0001 |
| MTphg | ~~ | MTphg | 1.2713 | 0.1813 | 0.3049 | 0.0472 | 6.4617 | <0.0001 |
| MDphg | ~~ | MDphg | 0.7379 | 0.1565 | 0.1681 | 0.0386 | 4.3505 | <0.0001 |
| Vpre | ~~ | Vpre | 1.0896 | 0.1919 | 0.2783 | 0.0516 | 5.3938 | <0.0001 |
| MTpre | ~~ | MTpre | 0.2797 | 0.1879 | 0.0647 | 0.0447 | 1.4461 | 0.1481 |
| MDpre | ~~ | MDpre | 0.5612 | 0.1281 | 0.1418 | 0.0334 | 4.2394 | <0.0001 |
| VLMT | ~~ | VLMT | 2.8932 | 0.1452 | 0.7241 | 0.0317 | 22.8525 | <0.0001 |
| FP | ~~ | FP | 3.1758 | 0.1418 | 0.7946 | 0.0276 | 28.8178 | <0.0001 |
| SE | ~~ | SE | 2.967 | 0.1438 | 0.7426 | 0.0306 | 24.2749 | <0.0001 |
| OL | ~~ | OL | 2.7044 | 0.1497 | 0.6764 | 0.0345 | 19.6222 | <0.0001 |
| **Latent variable** | **Variance** | | **Estimate** | **SE** | **Estimate std.** | **SE std.** | **Z** | **P-value** |
| PFC | ~~ | PFC | 0.3791 | 0.171 | 1 | 0 | NA | NA |
| HC | ~~ | HC | 1.7762 | 0.2627 | 1 | 0 | NA | NA |
| PHG | ~~ | PHG | 0.4598 | 0.1601 | 1 | 0 | NA | NA |
| PRE | ~~ | PRE | 1.2298 | 0.2156 | 1 | 0 | NA | NA |
| V | ~~ | V | 1.5965 | 0.2821 | 1 | 0 | NA | NA |
| MD | ~~ | MD | 1.9055 | 0.2843 | 1 | 0 | NA | NA |
| MT | ~~ | MT | 1.9239 | 0.3263 | 1 | 0 | NA | NA |
| **Latent variable** | **Residual variance** | | | | | | | |
| EM | ~~ | EM | 0.9602 | 0.1382 | 0.8711 | 0.0566 | 15.3972 | <0.0001 |

| Latent variable | Covariance | Latent variable | Estimate | SE | Estimate std. | SE std. | Z | P-value |
|---|---|---|---|---|---|---|---|---|
| PFC | ~~ | HC | 0.4817 | 0.1467 | 0.587 | 0.061 | 9.6168 | <0.0001 |
| PFC | ~~ | PHG | 0.2594 | 0.1002 | 0.6213 | 0.0751 | 8.2691 | <0.0001 |
| PFC | ~~ | PRE | -0.0538 | 0.0844 | -0.0788 | 0.1303 | -0.6049 | 0.5453 |
| HC | ~~ | PHG | 0.7428 | 0.178 | 0.822 | 0.0332 | 24.7337 | <0.0001 |
| HC | ~~ | PRE | 0.0353 | 0.1279 | 0.0239 | 0.0861 | 0.2772 | 0.7816 |
| PHG | ~~ | PRE | 0.1812 | 0.0806 | 0.241 | 0.087 | 2.7711 | 0.0056 |
| V | ~~ | MT | 0.8186 | 0.1964 | 0.4671 | 0.0761 | 6.1384 | <0.0001 |
| V | ~~ | MD | -1.1448 | 0.2067 | -0.6564 | 0.0518 | -12.6669 | <0.0001 |
| MT | ~~ | MD | -1.5627 | 0.2516 | -0.8162 | 0.0342 | -23.8639 | <0.0001 |
| **Observed variable** | **Residual covariance** | **Observed variable** | **Estimate** | **SE** | **Estimate std.** | **SE std.** | **Z** | **P-value** |
| Vmo | ~~ | Vdl | 0.6093 | 0.1987 | 0.2783 | 0.0723 | 3.8489 | 1.00E-04 |
| Vhc | ~~ | Vphg | 1.4039 | 0.1707 | 0.6296 | 0.0389 | 16.1732 | 0 |
| **Latent variable** | **Regression** | **Latent variable** | **Estimate** | **SE** | **Estimate std.** | **SE std.** | **Z** | **P-value** |
| EM | ~ | PFC | 0.108 | 0.2599 | 0.0633 | 0.1511 | 0.419 | 0.6752 |
| EM | ~ | HC | 0.1914 | 0.1532 | 0.243 | 0.1957 | 1.2414 | 0.2144 |
| EM | ~ | PHG | 0.1094 | 0.3833 | 0.0706 | 0.246 | 0.2871 | 0.7741 |
| EM | ~ | PRE | 0.0745 | 0.1075 | 0.0787 | 0.1132 | 0.6952 | 0.4869 |
| **Observed variable** | **Intercept** | | **Estimate** | **SE** | **Estimate std.** | **SE std.** | **Z** | **P-value** |
| Vmo | ~1 | | 4.9921 | 0.1099 | 2.498 | 0.1118 | 22.3526 | <0.0001 |
| MTmo | ~1 | | 4.9343 | 0.1298 | 2.4477 | 0.1371 | 17.8512 | <0.0001 |
| MDmo | ~1 | | 5.0246 | 0.1175 | 2.5107 | 0.1211 | 20.7388 | <0.0001 |
| Vdl | ~1 | | 4.9962 | 0.11 | 2.501 | 0.1119 | 22.354 | <0.0001 |
| MTdl | ~1 | | 4.9481 | 0.132 | 2.4487 | 0.1391 | 17.6049 | <0.0001 |
| MDdl | ~1 | | 5.1117 | 0.1183 | 2.5084 | 0.1179 | 21.2706 | <0.0001 |
| Vhc | ~1 | | 4.9763 | 0.109 | 2.4879 | 0.1091 | 22.7988 | <0.0001 |
| MThc | ~1 | | 4.9551 | 0.1158 | 2.4546 | 0.1222 | 20.0861 | <0.0001 |
| MDhc | ~1 | | 5.2121 | 0.1234 | 2.3543 | 0.1116 | 21.0993 | <0.0001 |
| Vphg | ~1 | | 4.9883 | 0.1089 | 2.515 | 0.1108 | 22.697 | <0.0001 |
| MTphg | ~1 | | 4.9037 | 0.1292 | 2.4013 | 0.1348 | 17.8079 | <0.0001 |
| MDphg | ~1 | | 5.167 | 0.1202 | 2.4662 | 0.1156 | 21.337 | <0.0001 |
| Vpre | ~1 | | 4.9944 | 0.109 | 2.5239 | 0.1091 | 23.144 | <0.0001 |
| MTpre | ~1 | | 4.96 | 0.1285 | 2.3855 | 0.1256 | 18.9952 | <0.0001 |
| MDpre | ~1 | | 5.0412 | 0.1145 | 2.5338 | 0.1166 | 21.7249 | <0.0001 |
| VLMT | ~1 | | 5.0015 | 0.0516 | 2.5022 | 0.0525 | 47.6818 | <0.0001 |
| FP | ~1 | | 5.0003 | 0.0517 | 2.5012 | 0.0525 | 47.6183 | <0.0001 |
| SE | ~1 | | 5.0015 | 0.0517 | 2.5021 | 0.0525 | 47.6474 | <0.0001 |
| OL | ~1 | | 4.9994 | 0.0515 | 2.5001 | 0.0524 | 47.7513 | <0.0001 |
| **Latent variable** | **mean** | | **Estimate** | **SE** | **Estimate std.** | **SE std.** | **Z** | **P-value** |
| PFC | ~1 | | 0 | 0 | 0 | 0 | NA | NA |
| HC | ~1 | | 0 | 0 | 0 | 0 | NA | NA |

| | | | | | | | |
|---|---|---|---|---|---|---|---|
| PHG | ~1 | 0 | 0 | 0 | 0 | NA | NA |
| PRE | ~1 | 0 | 0 | 0 | 0 | NA | NA |
| V | ~1 | 0 | 0 | 0 | 0 | NA | NA |
| MT | ~1 | 0 | 0 | 0 | 0 | NA | NA |
| MD | ~1 | 0 | 0 | 0 | 0 | NA | NA |
| EM | ~1 | 0 | 0 | 0 | 0 | NA | NA |

*Note. P-values from Wald test.*

### 3. Sex differences in variables

For sex differences in all variables of interest, see Table S1. Women outperformed men in verbal memory (VLMT), which could be expected (for a review, see Asperholm et al. 2019), but showed no advantage in the other memory tests. In many of the grey matter integrity variables, women showed values indicating higher integrity (higher prefrontal volumes, higher MT values, lower MD values). The literature on regional grey matter structural differences between sexes is so far mixed. For instance, a meta-analysis suggests the extent of differences in volume and density to depend on the brain region (Ruigrok et al. 2014). It is also not trivial to disentangle whether different ways of accounting for ICV might account for differences between studies in regional sex differences (Voevodskaya et al. 2014; Pintzka et al. 2015).

Table S4: Sex differences in all variables of interest

| Variable | Mean male | SD male | Mean female | SD female | P-value |
|---|---|---|---|---|---|
| VLMT | 8.2479 | 2.61713 | 8.7371 | 2.72146 | 0.0004 |
| FP | 0.2597 | 0.20769 | 0.273 | 0.21147 | 0.21844 |
| SE | 0.2724 | 0.14168 | 0.2825 | 0.13775 | 0.16199 |
| OL | 13.0787 | 4.17183 | 13.4245 | 3.8318 | 0.0946 |
| Vhc | 0.5554 | 0.0576 | 0.6103 | 0.05068 | <0.00001 |
| Vphg | 0.535 | 0.03865 | 0.557 | 0.04364 | <0.00001 |
| Vpre | 0.418 | 0.03865 | 0.4436 | 0.04027 | <0.00001 |
| Vmofc | 0.4522 | 0.04228 | 0.4782 | 0.04107 | <0.00001 |
| Vdlpfc | 0.3657 | 0.03623 | 0.3904 | 0.03215 | <0.00001 |
| MThc | 322.2943 | 34.58607 | 347.6178 | 22.48292 | <0.00001 |
| MTphg | 359.8157 | 17.34042 | 369.0587 | 11.78399 | 0.00001 |
| MTpre | 331.5453 | 19.50512 | 342.1202 | 17.07309 | 0.00006 |
| MTmo | 349.9700 | 17.11499 | 359.4151 | 15.66446 | 0.00007 |
| MTdl | 297.3529 | 17.56731 | 305.0321 | 27.22796 | 0.0263 |
| MDhc | 0.0014 | 0.00015 | 0.0013 | 0.00011 | <0.00001 |
| MDphg | 0.0014 | 0.00012 | 0.0013 | 8.00E-05 | <0.00001 |
| MDpre | 0.0012 | 0.00011 | 0.0012 | 0.00011 | <0.00001 |
| MDmo | 0.0012 | 0.00009 | 0.0012 | 0.00011 | 0.00327 |
| MDdl | 0.0012 | 0.00008 | 0.0011 | 0.00007 | <0.00001 |
| Age | 70.5278 | 4.10643 | 70.667 | 3.56062 | 0.48231 |
| Education | 14.6049 | 2.87097 | 13.7467 | 2.85068 | <0.00001 |

*Note. P-values from Welch T-test for unequal variances.*

### 4. Sex differences in association structures

Figure S1: First-order correlations among all variables by sex. Left: male participants, right: female participants.

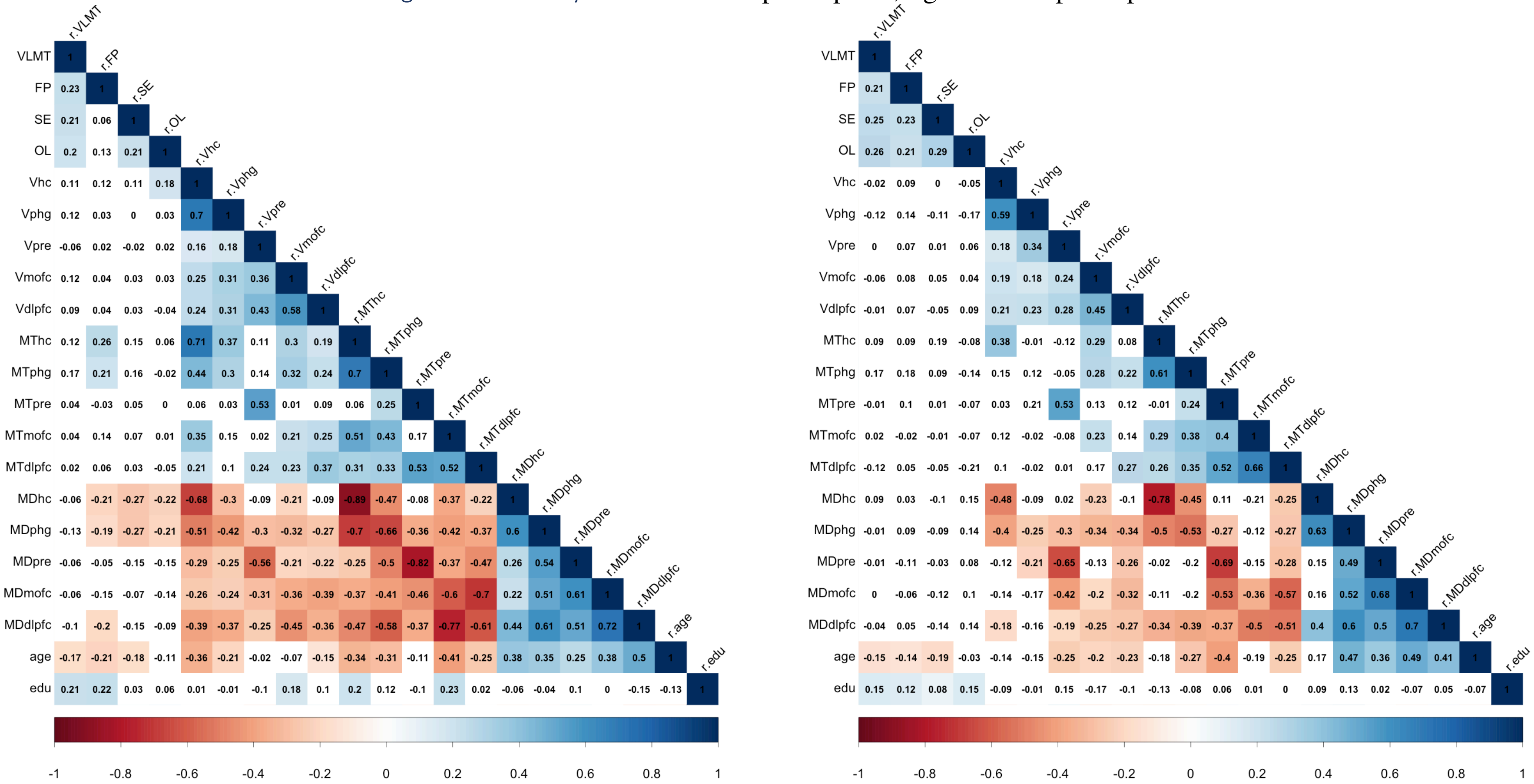

*Note*. *Color coding according to strength and direction of correlation (blue = positive, red = negative). White background: correlation non-significant (p >.05). VLMT = verbal learning and memory test; FP = face-profession task; SE = scene encoding; OL = object location task; V = VBM (volume-based morphometry); MT = Magnetization tranfer ratio; MD = mean diffusivity; hc = hippocampus; phg = parahippocampal gyrus; pre = precuneus; mofc = medio-orbitofrontal cortex; dlpfc = dorsolateral prefrontal cortex; edu = years of education.*

We tested the brain-only grey matter integrity MTMM for measurement invariance (Meredith 1993, Meredith and Teresi 2006) across sexes in the following way. First, we specified a two-group model in which we allowed all parameters to differ across groups (configural invariance). When we specified exactly the same model as for the total sample, a two-group model did not converge. A pruned version of the model, with the non-significant loadings on the PFC factor fixed to zero, namely that of MD in medio-orbitofronal cortex and MT in dorsolateral prefrontal cortex, fit the total sample data comparably well (Chi2difference (df = 2) = 3.56, $p$ = .17). This model was easier to fit to the two groups then. For fit indices of the two group model with configural invariance, that is, all parameters allowed to differ, see Table S4.

After fitting the configurally invariant model, we specified a model with all loadings restricted to be estimated at the same value for women and men (metric invariance). We compared the two models by their Chi2-values, checked the Chi2-difference and whether it is significant. Because the Chi2-difference depends on sample size and might be overly sensitive in large samples, we also checked the difference in CFI, RMSEA and SRMR for a relevant difference following rules-of-thumb based on a simulations study (Chen 2007) saying “when sample size is adequate (total N > 300) and sample sizes are equal across groups […], a CFI difference of -.01, supplemented by a change of >= .015 in RMSEA or a change of >= .03 in SRMR would indicate noninvariance” (Chen 2007). We then fit a model with loadings and residual variances of the indicator variables to be the same across sexes and compared it to the metric invariance model.

The Chi2 difference test and the RMSEA suggested a relevant worsening of fit when assuming invariant residuals as well, as compared to metric invariance, but the changes in RMSEA and SRMR did not reach the cutoffs, so that we continued assuming invariance of residuals as well (Table S5).

Table S5: Measurement invariance tests for the brain-only MTMM-model

| Invariance level | Df | AIC | BIC | Chi2 | Chi2diff | Pr(>Chi2) | CFI | RMSEA | SRMR |
|---|---|---|---|---|---|---|---|---|---|
| same model (configural) | 132 | 14331 | 14856 | 221 | | | 0.962 | 0.0638 | 0.0691 |
| same loadings (metric) | 153 | 14321 | 14765 | 252 | 31.4 | 0.0680 | 0.958 | 0.0626 | 0.0805 |
| same loadings and residuals | 168 | 14334 | 14722 | 295 | 43.5 | 0.0001 | 0.945 | 0.0678 | 0.1030 |

*Note. Df: degrees of freedom; AIC: Akaike information criterion; CFI: Comparitive fit index; RMSEA: root mean squared error of approximation; SRMR: standardized root mean square residual*

For episodic memory measurement invariance, see Table S6. Here, the above cutoffs suggested invariance of loadings and  of residuals.

Table S6: Measurement invariance tests for the episodic memory measurement model

| Invariance level | Df | AIC | BIC | Chisq | Chisq diff | Df diff | Pr(>Chi2) | CFI | RMSEA | SRMR |
|---|---|---|---|---|---|---|---|---|---|---|
| same model (configural) | 4 | 24822.6 | 24950.4 | 1.0456 | | | | 1.0000 | 0.0000 | 0.0052 |
| same loadings (metric) | 7 | 24819.3 | 24931.2 | 3.7634 | 2.7178 | 3 | 0.4372 | 1.0000 | 0.0000 | 0.0139 |
| same loadings and residuals | 11 | 24819.8 | 24910.3 | 12.2636 | 8.5002 | 4 | 0.0749 | 0.9974 | 0.0123 | 0.0276 |

*Note. Df: degrees of freedom; AIC: Akaike information criterion; CFI: Comparitive fit index; RMSEA: root mean squared error of approximation; SRMR: standardized root mean square residual*

We fit a multigroup model that includes both the grey matter model and episodic memory, with loadings and residuals fixed to be equal across sexes in the brain model. From this model we then obtained the estimated correlations between episodic memory and each of the grey matter ROIs. Please see Table S7 for the results. The results suggest that there are significant sex differences in the association between hippocampus and episodic memory as well as between parahippocampal gyrus and episodic memory. The associations found in the total sample are restricted to men. When adding age and education into the model (Table S8), the pattern stays the same, only that these associations are slightly attenuated within the male group.

Table S7: Associations between episodic memory and each of the ROI-wise integrity factors by sex, from a multigroup model

| Association of episodic memory with | Men covariance (SE) | Men correlation / std. cov | Women covariance (SE) | Women correlation / std. cov | Chi2 difference when setting covariance equal (df =1) | P-Value from Chi2 difference |
|---|---|---|---|---|---|---|
| PFC | .20 (.10) | .24 | -0.004(.09) | -.008 | 2.4 | .12 |
| HC | .58 (.10) | .39 | -.042 (.13) | -.02 | 8.97 | .0027 |
| PHG | .27 (.10) | .32 | -0.04 (.08) | -.09 | 6.54 | .011 |
| PRE | .08 (.14) | .07 | 0.10 (.17) | .09 | 0.00516 | .94 |

*Note. PFC: prefrontal cortex; HC: hippocampus; PHG: parahippocampal gyrus; PRE: precuneus*

Table S8: Associations between episodic memory and each of the ROI-wise integrity factors by sex, from a multigroup model with age and education as covariates

| Association of episodic memory with | Men covariance (SE) | Men correlation / std. cov | Women covariance (SE) | Women correlation / std. cov | Chi2 difference when setting covariance equal (df =1) | P-Value from Chi2 difference |
|---|---|---|---|---|---|---|
| PFC | .02 (.08) | .03 | -0.03 (.09) | -.05 | 0.154 | .70 |
| HC | .37 (.14) | .29 | -0.04 (.13) | -.05 | 4.76 | .029 |
| PHG | .18 (.09) | .24 | -0.06 (.07) | -.16 | 4.5 | .034 |
| PRE | .09 (.13) | .08 | 0.03 (.17) | .03 | 0.067 | .8 |

*Note. PFC: prefrontal cortex; HC: hippocampus; PHG: parahippocampal gyrus; PRE: precuneus*

As this is the result of an exploratory, additional analysis, we can only speculate post-hoc about the reasons for the associations to be restricted to men. We do not believe that fundamentally different processes are at work when it comes to brain integrity and cognitive aging in men and women; therefore, we did not expect these differences and did not consider them when conceptualizing this study. Rather, a larger share of the men than of the women might already have proceeded further on their brain aging trajectory, are beginning to suffer cognitive consequences. To speculate, in our male population, metabolic risk might be more prevalent, with detrimental effects on both grey matter integrity (Raz et al. 2005; Raz and

Rodrigue 2006) and episodic memory (Yates et al. 2012). This might also be the cause of men showing lower mean brain integrity estimates and lower mean episodic memory performance than women in the present study (Tables 3 and S2). The fact that we see large interindividual variance in both males and females is not contradicting this idea, given that interindividual differences have probably to the largest extent been there throughout the lifespan and are not distinguishable from aging-related changes in this cross-sectional data set. Future studies including longitudinal follow-ups and measures of metabolic risk may shed light on these questions.
